# Convergent flow-mediated mesenchymal force drives embryonic foregut constriction and splitting

**DOI:** 10.1101/2025.01.22.634318

**Authors:** Rui Yan, Ludwig A. Hoffmann, Panagiotis Oikonomou, Deng Li, ChangHee Lee, Hasreet K. Gill, Alessandro Mongera, Nandan L. Nerurkar, L. Mahadevan, Clifford J. Tabin

## Abstract

The transformation of a two-dimensional epithelial sheet into various three-dimensional structures is a critical process in generating the diversity of animal forms. Previous studies of epithelial folding have revealed diverse mechanisms driven by epithelium-intrinsic or -extrinsic forces. Yet little is known about the biomechanical basis of epithelial splitting, which involves extreme folding and eventually a topological transition breaking the epithelial tube. Here, we leverage tracheal-esophageal separation (TES), a critical and highly conserved morphogenetic event during tetrapod embryogenesis, as a model system for interrogating epithelial tube splitting both in vivo and ex vivo. Comparing TES in chick and mouse embryos, we identified an evolutionarily conserved, compressive force exerted by the mesenchyme surrounding the epithelium, as being necessary to drive epithelial constriction and splitting. The compressive force is mediated by localized convergent flow of mesenchymal cells towards the epithelium. We further found that Sonic hedgehog (SHH) secreted by the epithelium functions as an attractive cue for mesenchymal cells. Removal of the mesenchyme, inhibition of cell migration, or loss of SHH signaling all abrogate TES, which can be rescued by externally applied pressure. These results unveil the biomechanical basis of epithelial splitting and suggest plausible mesenchymal origins of tracheal-esophageal birth defects.

## Introduction

The epithelium, a fundamental tissue type across all metazoans, can adopt morphologies ranging from simple, two-dimensional sheets to highly complex, three-dimensional scaffolds, giving rise to the characteristic forms and thus functions of different organs. How the epithelium folds during morphogenesis of these organs has been a central question in mechanobiology^1–3^. We now know that epithelial folding can be driven by both intrinsic and extrinsic mechanisms. Epithelium-intrinsic forces such as those generated by apical/basal constriction or directional cell migration, are crucial for many epithelial morphogenetic events including gastrulation and neural tube closure^1,3,4^. Conversely, non-epithelial tissue can also generate force to shape the adjacent epithelium, for example, through differential proliferation or cell clustering^2,5–7^.

Despite these growing insights into epithelial folding, very little is known about the extreme case in which epithelial tube deformation reaches a state where closely apposed folded segments are reconfigured to form a new junction. A transient junction may relax without changing the topology of the epithelium, whereas a sustained junction can potentially lead to reorganization of epithelial cell polarity, resulting in a topological transition that breaks the continuity of the epithelial tube, which we here define as epithelial splitting. This is seen in tracheal-esophageal separation/septation (TES) in tetrapods, as well as cloacal septation in mammals, in both cases involving the splitting of an epithelial tube into two tubes giving rise to different organs^8–10^. Although the morphological and genetic aspects of TES have been extensively studied in the past two decades, very few studies have examined the biomechanical basis of epithelial splitting in this intriguing system, and the biophysical force drive the formation and resolution of the epithelial junction remain unclear^11^.

TES starts with the dorsal-ventral patterning of the foregut epithelium into esophageal and tracheal progenitors at Embryonic Day 9.5 (E9.5) in mouse embryos, mainly attributed to ventral-to-dorsal gradients of BMP and Wnt signaling and Ephrin-mediated cell sorting^8,9,12,13^. Over the next two days, the foregut epithelium constricts bilaterally and forms a medial junction (septum) where cells transiently lose and re-establish their apicobasal polarity, resolving the septum to generate the esophagus dorsally and the trachea ventrally (Fig. 1a) ^14,15^. TES is highly conserved in tetrapods, and defects in TES cause a spectrum of congenital tracheal-esophageal malformations impacting 1 in 2,500 humans^14–16^. Using mouse and frog embryos as models, previous studies have identified a number of genetic factors essential for the epithelial patterning, sorting, and septal resolution^11,13–15,17–21^. Yet the driving force of epithelial constriction remains unknown. So far, there are more than 50 genes associated with tracheal-esophageal birth defects in humans and ∼15 genes whose loss-of-function mutations lead to defects in mouse TES^14^. Among these genes, some are expressed in the epithelium, whereas others are expressed in the surrounding mesenchyme, a subset of which do not have an obvious connection to the morphogen gradients patterning the epithelium. It remains an open question whether TES is driven by epithelium-intrinsic forces, or requires additional force from the mesenchyme to achieve its extreme folded geometry and to enable splitting. Addressing this question will not only help explain the etiology of TES-related birth defects, but also deepen our understanding of the biophysical mechanism driving epithelial splitting.

**Figure 1.**
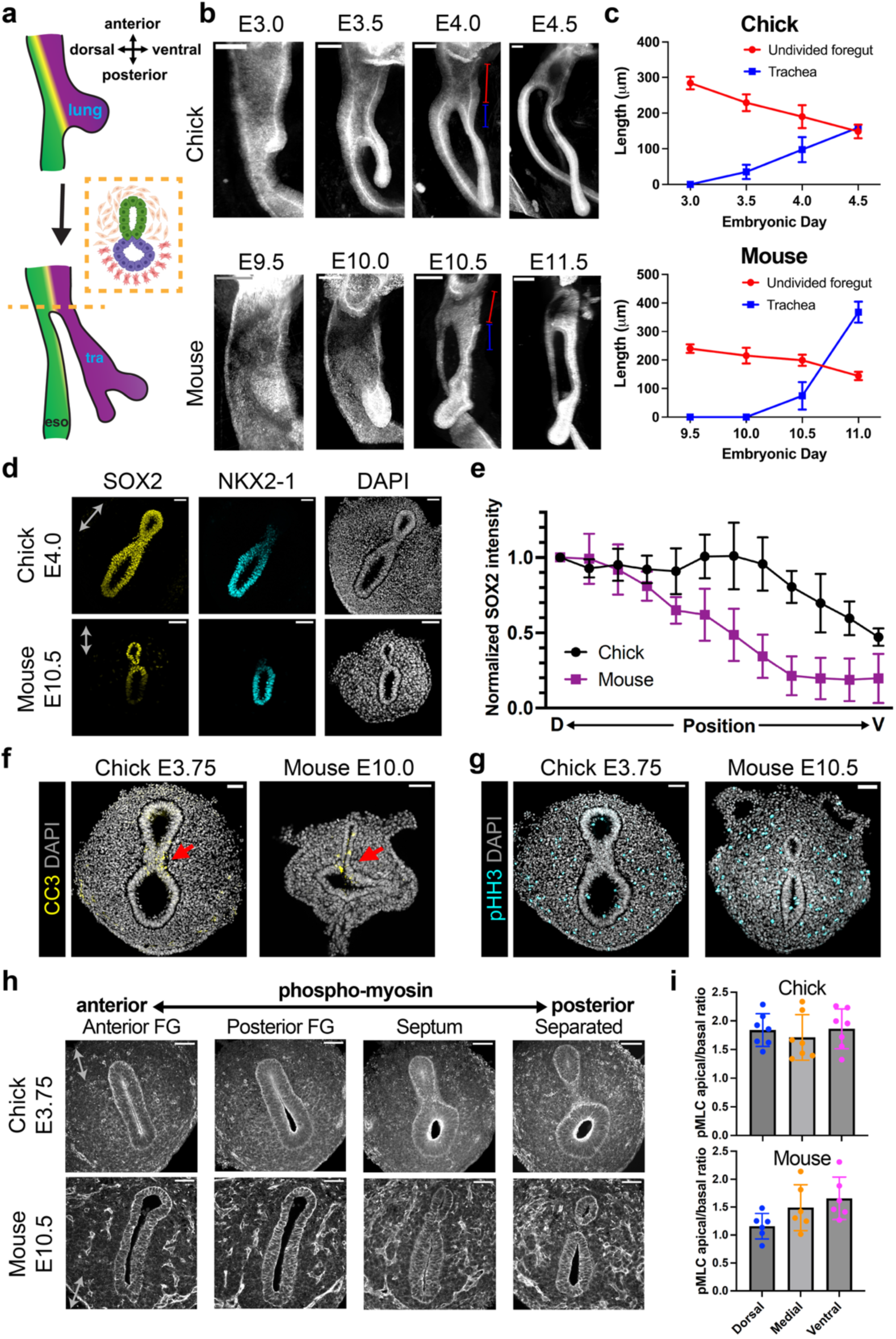
Morphological and molecular comparison of chick and mouse tracheal-esophageal separation (TES), a model system for studying epithelial splitting. **(a)** Schematic of TES. The airway (purple) and the esophagus (green) form through foregut splitting. The inset shows the cross-section of the foregut at the level of TES along the dashed line. **(b)** Whole mount immunofluorescence of E-cadherin (CDH1) of chick and mouse foreguts during TES. Images show lateral views from the right. The red and blue line segments indicate the lengths of the undivided foregut and the trachea. **(c)** Quantification of the lengths of the undivided foregut (red) and the trachea (blue) during TES from whole mount images. N = 4 biological replicates for each stage. **(d)** Immunofluorescence of SOX2 and NKX2-1 in transverse sections of the chick and mouse foreguts during TES. **(e)** Quantification of SOX2 intensity along the dorsal-ventral axis (N = 6 samples). **(f,g)** Immunofluorescence of cleaved Caspase 3 (CC3) (f) and phospho-Histone H3 (pHH3) (g) in transverse sections of chick and mouse TES. Arrows point to the tracheal-esophageal septum. **(h)** Immunofluorescence of phospho-MLC2 (pMLC) in serial transverse sections of chick and mouse foreguts, from the anterior undivided foregut through the posterior separated foregut. **(i)** Quantification of the apical-basal ratio of pMLC intensity in the dorsal, medial, and ventral foregut epithelium (N = 6 to 7 samples for each species). Scale bars: 100 µm (b), 50 µm (d,f-h). Error bars indicate standard deviation (SD). Double arrows indicate the dorsal-ventral axes for tilted samples.

To this end, we comparatively studied TES in the chick and the mouse to identify evolutionarily conserved biophysical mechanisms of epithelial splitting. Far less prior work has been done on chick TES^8^. We found it to be morphologically similar to mouse TES, making it plausible to extract evolutionarily conserved mechanisms from the two systems (Fig. 1b). To enable access to biophysical force measurements, we focused on analyzing live tissues, and we established an ex vivo slice culture system in which TES can be visualized in real-time, pharmacologically perturbed, and surgically manipulated. We found that although chick and mouse TES differ in many aspects such as epithelial morphology and cytoskeletal structure, both species rely on forces generated by the mesenchyme to facilitate epithelial constriction, which itself is also sufficient to induce septation. We further discovered that the mesenchymal cells flow convergently towards the epithelium causing the compressive stresses in both species. Sonic hedgehog (SHH) secreted by the epithelium endogenously attracts surrounding mesenchymal cells and modulates the epithelium’s shape. These results demonstrate the essential role of mesenchymal force in epithelial splitting and suggest that defects in mesenchymal cell migration in response to SHH signaling is a plausible mechanism of tracheal-esophageal malformations.

## Results

### Morphological and molecular comparison of chick and mouse TES

We first thoroughly characterized chick TES in vivo, such that it can be spatiotemporally aligned to the better-studied mouse system. Chick lung buds emerge at ∼E3.0, or Hamburger Hamilton (HH) stage 17-18, and then elongate posteriorly. TES initiates shortly after at E3.5 (HH19), from the lung budding site moving anteriorly, until the TES septum reaches the larynx at E4.5 (HH23-24) (Fig. 1a,b, Supplementary Movie 1). The undivided anterior foregut shortens by ∼150 µm in 1.5 days, a length equal to the nascent trachea (Fig. 1c). A similar rate of foregut shortening is observed in mouse TES, whereas the mouse trachea exhibits additional elongation (Fig. 1b,c, Supplementary Movie 2). The foregut epithelia in both species are pseudostratified (Supplementary Fig. 1a). The foregut morphology is dorsoventrally asymmetrical because of the gradients of BMP and Wnt signaling. Molecularly, the ventral foregut (prospective trachea) of the chick embryo is marked by NKX2-1, and the dorsal foregut (prospective esophagus) is marked by SOX2 (Fig. 1d), markers previously described in the context of mouse TES^17,22^. Of note, the dorsal-ventral gradient of chick SOX2 is less pronounced than the mouse (Fig. 1e), implying that NKX2-1 and SOX2 may not be mutual inhibitory in the chick as they have been reported to be in the mouse^17,19,23^. At the septum, many epithelial cells undergo apoptosis, which is rarely observed outside this region (Fig. 1f). Cell proliferation is also halted at the septum in both species, whereas it appears homogenous across the mesenchyme (Fig. 1g), consistent with what has been reported^11,24^.

We further probed the molecular patterns pertinent to tissue biomechanics, including localization of phosphorylated myosin light chain (p-MLC), F-actin, and extracellular matrix components. In the chick, p-MLC and F-actin are highly enriched on the apical side of the epithelium outside the septum, where cells lose their polarity (Fig. 1h, Supplementary Fig. 1b). This pattern suggests strong apical constriction in the chick epithelium. By contrast, p-MLC is not highly enriched apically in the mouse (Fig. 1h), despite an apical concentration of actin (Supplementary Fig. 1b). The ratio of apical to basal p-MLC, an indirect measure of apical constriction, indicates a weaker epithelium-intrinsic constrictive force in the mouse than in the chick (Fig. 1i). Electroporation of tdTomato-tagged MLC2 into the chick foregut epithelium revealed a less elongated epithelial morphology in cells undergoing tube splitting compared to the cells outside this region (Supplementary Fig. 1c,d). Hyaluronic acid, which is known for generating fluid pressure within the tissue^25^, shows a dorsal-ventral gradient in the chick but a homogeneous distribution in the mouse (Supplementary Fig. 1e). Fibronectin, a stiffer component of the extracellular matrix, is ubiquitously expressed in the mesenchyme in both species (Supplementary Fig. 1f), These results suggest that although chick and mouse TES are morphologically similar, they exhibit significant differences related to tissue biomechanics at the molecular and cellular scales. It thus became imperative to examine the tissue-level biomechanics to understand this conserved morphogenetic program.

### Evolutionarily conserved, constrictive mesenchymal force deforms the epithelium during TES

We cut freshly dissected foreguts transversely into ∼100 µm-thick slices to obtain a cross-sectional view of epithelial splitting in live tissue (Fig. 2a). To probe the force distribution at the epithelium, we surgically removed the surrounding mesenchyme and analyzed the epithelial deformation (Fig. 2a). Intriguingly, the epithelium expanded immediately after mesenchymal removal, indicating that the epithelium was being compressed by the mesenchyme (Fig. 2a, Supplementary Fig. 2a,b). The expansion was more pronounced mediolaterally than dorsoventrally. The results were consistent in the chick and the mouse, and a similar pattern of epithelial deformation was previously reported in *Xenopus* foregut^11^, implying that such constrictive mesenchymal force is evolutionarily conserved.

**Figure 2.**
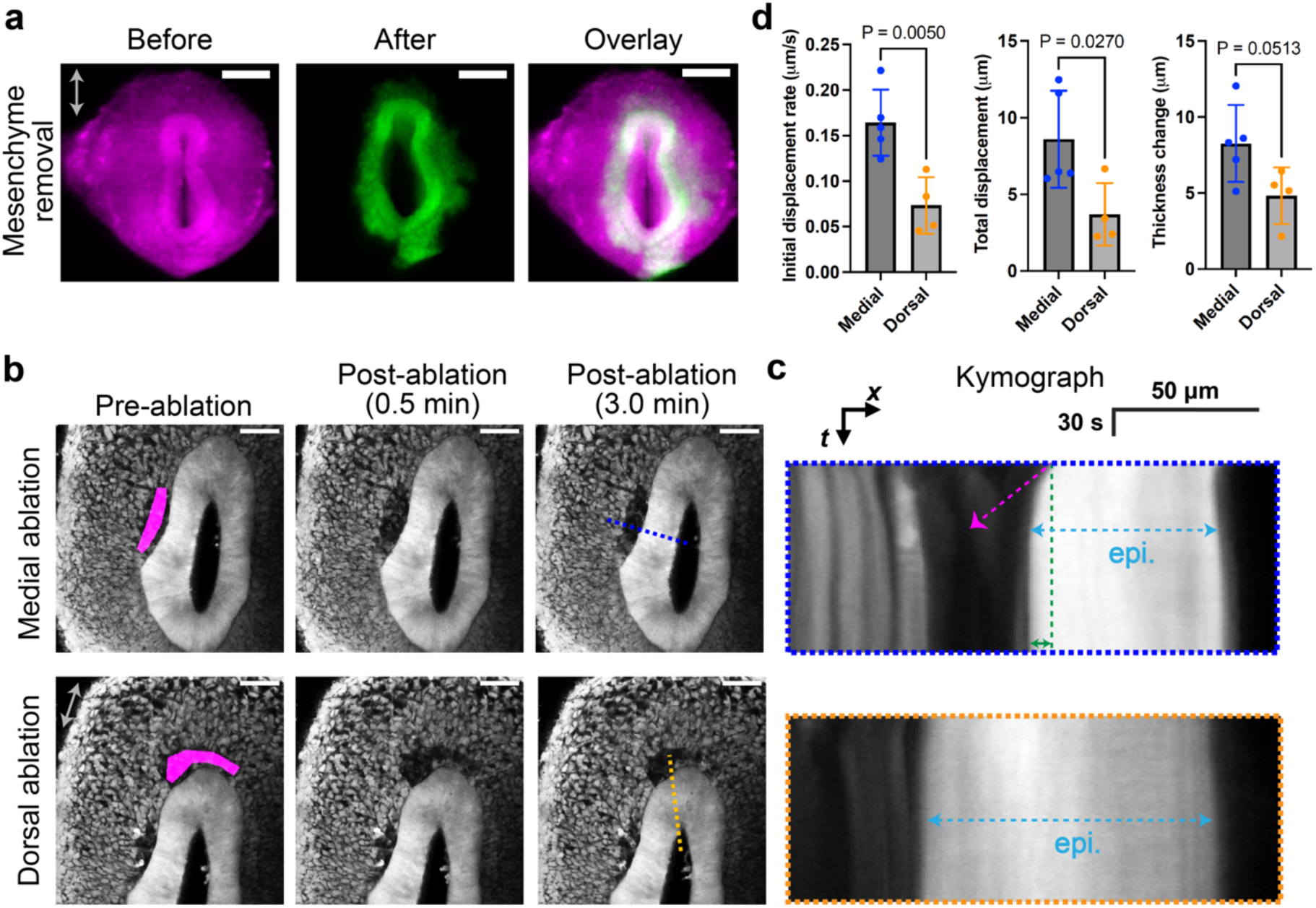
Constrictive mesenchymal force deforms the epithelium during TES. **(a)** Stereoscope imaging of the transverse slice of an E3.75 GFP chick foregut before (magenta) and after (green) surgical removal of the mesenchyme. **(b)** Two-photon laser ablation of an E3.75 GFP chick foregut slice in the medial or dorsal sub-epithelial mesenchyme (magenta). **(c)** Kymographs along the dotted lines in (b). Magenta arrowhead indicates the initial displacement rate. The green line and double arrows indicate the total epithelial displacement. The light blue dashed double arrows mark the epithelial thickness. **(d)** Quantifications of dorsal and medial ablation measurements as in (c). N = 4 biological replicates. Scale bars: 100 µm (a), 50 µm (b). Error bars: SD. P values: Two-tailed Welch’s t test.

We further confirmed this finding with localized laser ablation of the sub-epithelial mesenchyme. When a fraction of the medial mesenchyme was ablated, the epithelium moved towards the ablated site, reminiscent of the bulk tissue removal assay (Fig. 2b,c, Supplementary Fig. 2b,c, Supplementary Movie 3-5). By comparison, no significant epithelial deformation was observed when the dorsal mesenchyme was ablated (Fig. 2b-d, Supplementary Fig. 2b-d). Quantitative analyses of the initial displacement rate and total displacement suggest that a compressive mesenchymal force exists near the medial region of the epithelium, whose loss leads to both lateral movement and thickening of the epithelium (Fig. 2d, Supplementary Fig. 2d). Altogether, we conclude that the mediolateral constriction by the mesenchyme is an evolutionarily conserved feature of TES.

### The mesenchymal force is essential for TES ex vivo

Having demonstrated the presence of a constrictive mesenchymal force, we next asked whether this force is functionally important to TES morphogenesis. This requires a system allowing real-time monitoring of TES as well as experimental perturbations. To that end, we developed an ex vivo culture of foregut slices in Matrigel, which, for at least 24 hours, preserved the dorsoventral patterning of SOX2 and NKX2-1 and the spatial distribution of cell proliferation (Fig. 3a, Supplementary Fig. 3a,b). There was detectable, but insubstantial apoptosis in the ex vivo culture (Supplementary Fig. 3c). Importantly, the slice culture recapitulates TES morphogenesis in vivo for both chick and mouse, which could be visualized by two-photon live imaging (Fig. 3b, Supplementary Movie 6,7). The observed morphological changes were not due to the focusing plane shift during acquisition, as we could track individual 200-nm fluorescent beads embedded in this system for >12 hours (Supplementary Fig. 3d,e, Supplementary Movie 8), and consistent morphological changes were observed across different imaging planes (Supplementary Fig. 3f). The decay of image intensity over long periods of imaging appears to be due to photobleaching (Supplementary Fig. 3g). To quantitatively characterize the morphological parameters of the epithelium, we used a machine learning-based automated segmentation pipeline to extract epithelial morphology from the time-lapse imaging data (Supplementary Movie 6, see also Methods). The epithelium narrows to create negative curvatures in the medial region. The distance between the bilateral curvature minima, defined as the neck width, continuously decreases during TES, whereas the area of the epithelium exhibits minimal changes (Fig. 3a,c,d). The dorsoventral length of the epithelium thus increases, and we use the dimensionless ratio of the epithelial perimeter to the square root of the area (perimeter-area ratio) to indicate the elongation of the epithelium (Fig. 3e). As the epithelium narrows during TES, the neck curvature also decreases monotonically (Fig. 3f). Although TES at different time points have distinct epithelial morphologies (Fig. 3b, Supplementary Fig. 4), the shape evolution in the neck width and the perimeter-area ratio is highly similar, especially after the formation of the septum (Fig. 3c,e).

**Figure 3.**
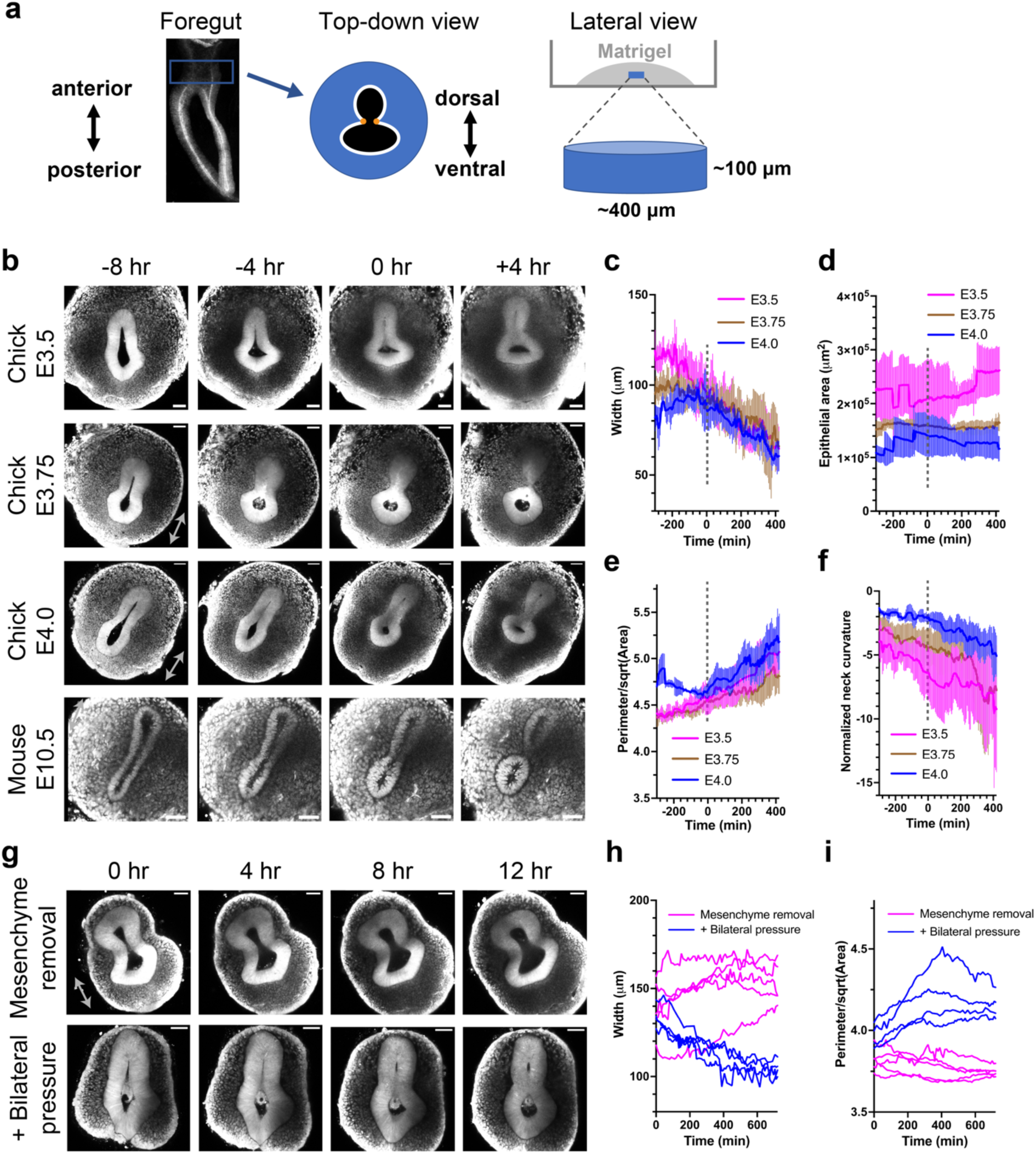
Mesenchymal force is essential for TES in ex vivo foregut slice culture. **(a)** Schematics of foregut slice culture. The white contour depicts the perimeter of the epithelium, the black shade indicates the area enclosed by the epithelium, and the distance between the two orange dots defines the neck width. **(b)** Live imaging of GFP chick and nTnG mouse foregut slices in culture. The moment of formation of the epithelial septum is defined as time 0. **(c-f)** Quantification of the neck width (c, see also Methods), the area enclosed by the epithelium (d), the ratio between the epithelial perimeter and the square root of its area (e, referred to as the perimeter-area ratio below), and the normalized neck curvature of the epithelium (f) from chick slice culture videos. Data are shown as median ± median absolute deviation (MAD). N = 9-12 biological replicates for each time point. The dotted lines indicate time 0 when the epithelial septum forms, which is used to align different samples temporally. **(g)** Live imaging of E3.75 GFP chick slices with surgical removal of the mesenchyme, in the absence (top) or the presence (bottom) of bilateral pressure by a collagen gel trough. **(h,i)** Quantification of the neck width (h) and the perimeter-area ratio (i) of mesenchyme-removed slices without (magenta) or with external bilateral force (blue). Each curve shows an individual sample (N = 5 for slices with only mesenchyme removal, N = 4 for slices with mesenchyme removal and external pressure). Scale bars: 50 µm.

Leveraging the slice culture system, we then asked how the absence of the mesenchyme would affect TES. Strikingly, mesenchyme-reduced foregut slices failed to complete TES (Fig. 3g). Instead, their initial constriction in the epithelium often reversed over time, even for those with an existing septum (Fig. 3g, Supplementary Movie 9,10). This suggests the constrictive mesenchymal force may be essential for TES ex vivo. However, an alternative explanation could be that the reduction of the mesenchyme also diminished the mesenchymal signaling molecules which are potentially important to TES. To distinguish between these possibilities, we squeezed mesenchyme-removed slices into a narrow trough made of a stiff collagen gel, which provides external forces in the mediolateral direction as the remaining mesenchyme proliferates in such bilaterally confined space. Such external compression rescued the morphological progression of TES (Fig. 3g-i, Supplementary Fig. 5, Supplementary Movie 11), indicating an essential biomechanical role of the foregut mesenchyme in epithelial splitting during TES.

### Directed cell migration underlies the essential compressive mesenchymal force

We next investigated the cellular basis of the mesenchymal force. It is well understood that local compressive force can be generated by differential proliferation within the tissue^5,26,27^. We thus performed pulsed EdU labeling to test this possibility. We found that the medial mesenchymal cells do not proliferate at a faster rate than other parts of the mesenchyme (Fig. 4a,b), in line with previous findings in the mouse using proliferation markers^11,24^, and thus cannot explain the mediolateral enrichment and the anisotropy of mesenchymal force. Moreover, when we inhibited cell proliferation with aphidicolin in the slice culture, the number of mitotic cells decreased (Fig. 4c), but the morphological features of TES were not affected (Fig. 4d-f, Supplementary Movie 12). Another possibility that mesenchymal cells undergo active elongation to generate pressure was not supported at the population level, even though some cells appeared elongated as they invaded the basement membrane (Fig. 3b, Supplementary Fig. 6).

**Figure 4.**
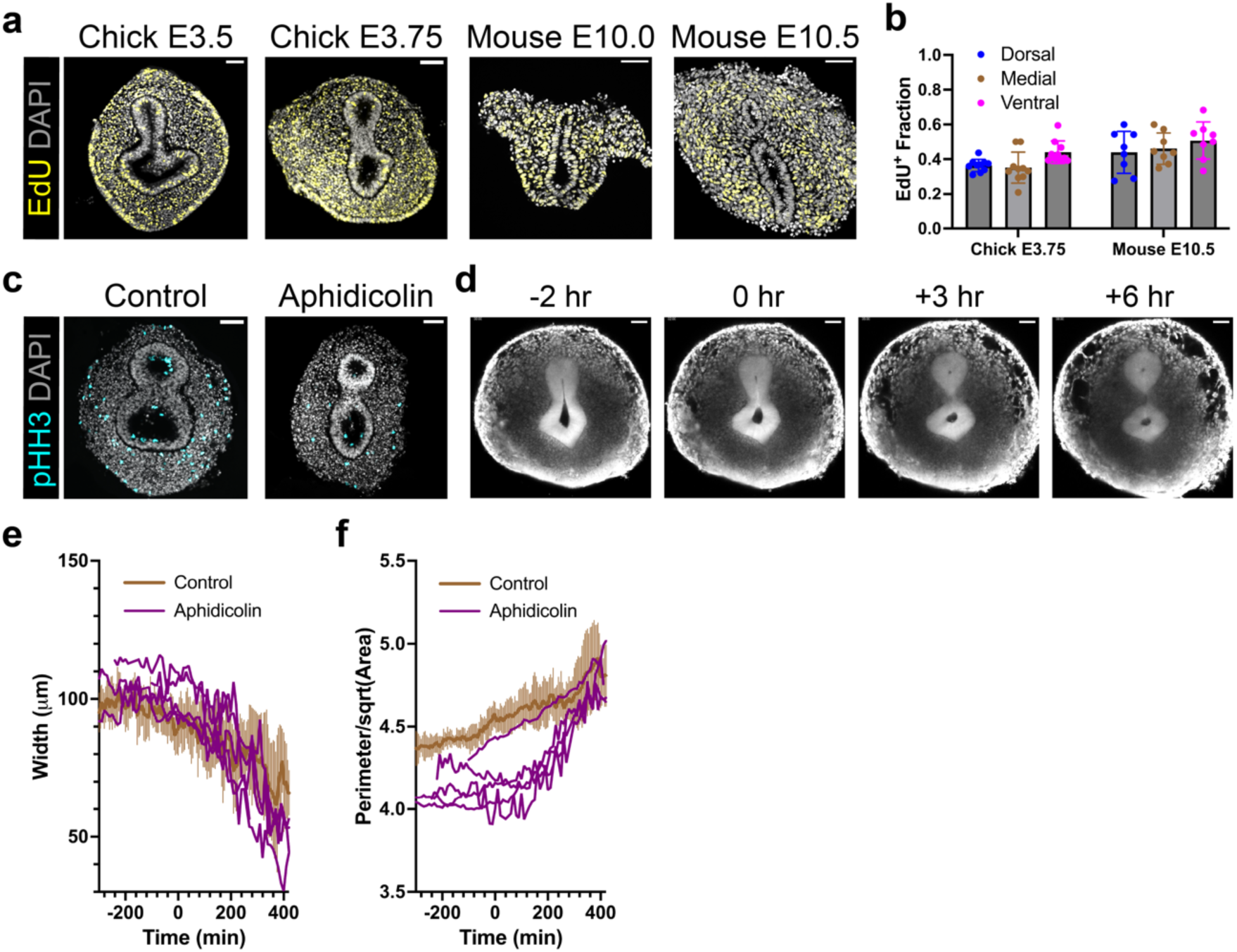
Cell proliferation does not explain the mesenchymal force distribution and is dispensable for TES. **(a)** Fluorescence imaging of transverse sections of chick and mouse slices ex vivo fixed after 10 µM EdU labeling for 2 hours. **(b)** Quantification of EdU-positive cells in the mesenchyme along the dorsoventral axis. Data are presented as mean ± SD. Each dot represents one slice. **(c)** Immunostaining of phospho-Histone H3 in E3.75 chick slices cultured with or without 3 µM aphidicolin for 24 hours. **(d)** Live imaging of an E3.5 GFP chick slice treated with 3 µM aphidicolin. **(e,f)** Quantification of the neck width (e) and the perimeter-area ratio (f) of aphidicolin-treated E3.75 chick slices (purple) overlaid with E3.75 controls (brown, as in Fig. 3c,e). Each curve shows an individual sample (N = 5). Scale bars: 50 µm.

An alternative source of localized force could be directional cell flow^28^. We hypothesized that if the mesenchymal cells converged towards the epithelium, they could generate a mediolateral pressure potentially driving TES. Harnessing the single-cell resolution in our live imaging of the slice culture, we performed particle image velocimetry (PIV) analysis to infer the temporally resolved velocity field of mesenchymal cells (Fig. 5a, Supplementary Movie S13, see Methods). This revealed that the mesenchymal cells indeed converged towards the epithelium, and the tissue-level strain inferred from the velocity field is initiated in the mesenchyme which then propagates to the epithelium (Fig. 5a, Supplementary Fig. 7a,b). As the chick mesenchymal cells are more densely packed than the mouse, the source of tissue strain appeared less pronounced, but the velocity field shows the same trend (Supplementary Fig. 7c). Focusing on the mouse, we independently validated the convergent mesenchymal motion by single-particle tracking of nuclear-labeled cells (Fig. 5b, Supplementary Movie 14). Analysis of the tracked cells’ trajectories showed that the cells converged towards the epithelium with varied average velocities yet strongly biased angle distribution (Fig. 5c-e). Mean squared displacement (MSD) analysis further revealed an apparent super-diffusion of cells with a scaling exponent of ∼1.2, suggesting active cell migration (Fig. 5f). To rule out the potential artifacts by analyzing the three-dimensional (3D) tissue in 2D, we also performed cell segmentation and tracking in 3D, which yielded migration patterns consistent with 2D analysis (Supplementary Fig. 8, Supplementary Movie 15). However, as many datasets were not suitable for 3D analysis due to the deterioration of image contrast in deeper tissue, we performed most analyses in 2D.

**Figure 5.**
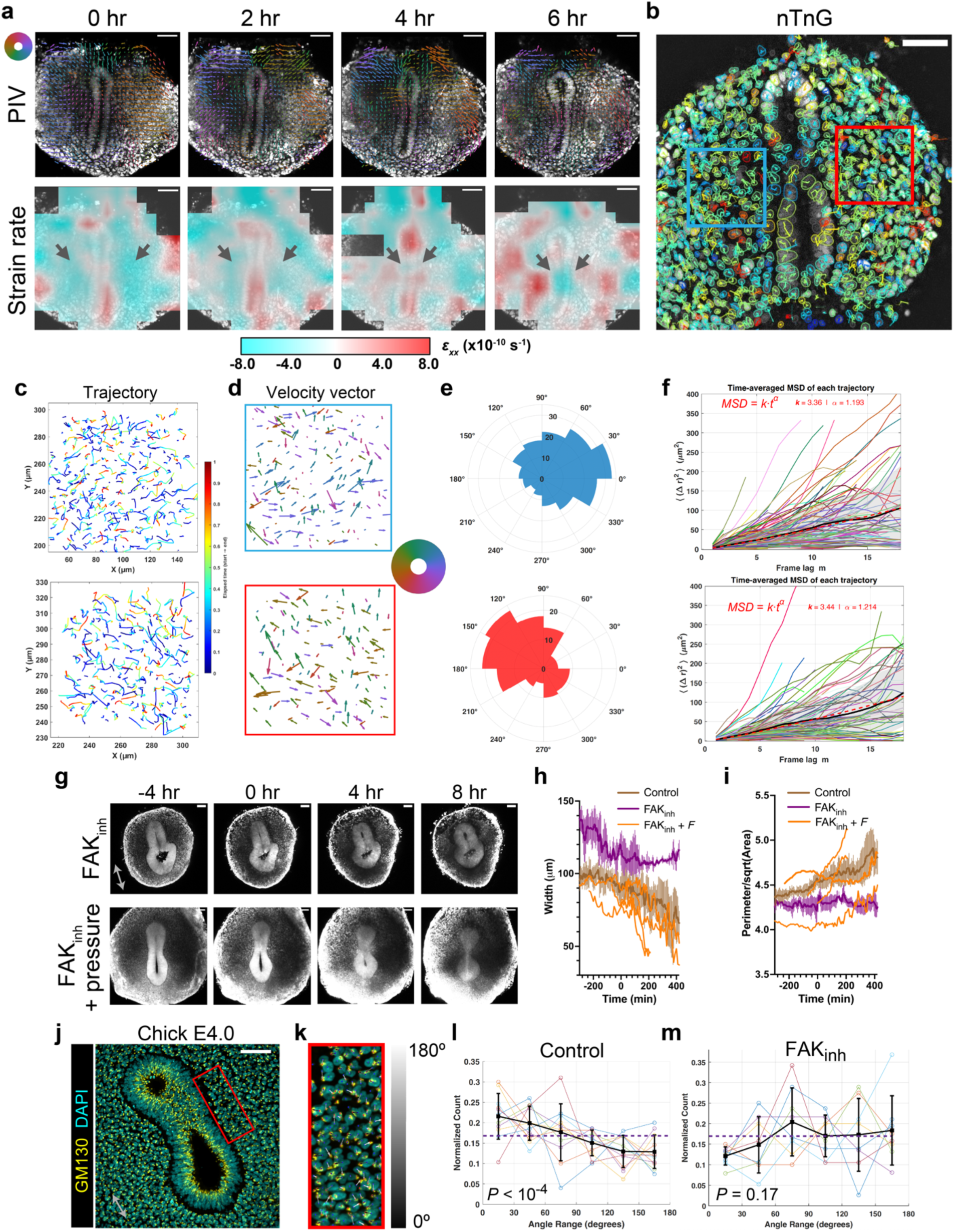
Directional mesenchymal flow contributes to the convergent mesenchymal force and is essential for TES. **(a)** Particle image velocimetry (PIV) analysis of an E10.5 nTnG mouse foregut slice culture and mapping of the strain rate along the mediolateral axis (*ε_xx_*). Cyan indicates compressive strain and red indicates expansive strain. Arrows indicate the shift of the compressive strain from the mesenchyme to the epithelium. The color wheel indicates the directions of PIV vectors. **(b)** Live imaging of an E10.5 nTnG mouse slice overlaid with cellular trajectories colored by their average speeds. **(c)** Individual cell trajectories in the blue (upper) and red (lower) boxes in (b). Trajectories are color-coded by time. **(d)** Average velocity vectors of individual cell trajectories in (c). Vectors are scaled with the speed and color-coded by their directions according to the color wheel. **(e)** Angular histograms the velocity vectors in (d). **(f)** Mean squared displacement (MSD) plots of cell trajectories in the blue (upper) and red (lower) boxes in (b). The black lines and shades show the population averages and standard deviations. The red dashed lines indicate the power-law fitting results. Frame time: 7.5 minutes. **(g)** Live imaging of E3.75 GFP chick slices with 2.5 µM FAK inhibitor (PF-573228), in the absence (top) or the presence (bottom) of bilateral pressure by a collagen gel trough. **(h,i)** Quantification of the neck width (h) and the perimeter-area ratio (i) of FAK inhibitor-treated slices without (purple) or with external bilateral force (orange) compared with E3.75 controls (brown, as in Fig. 3c,e). Averaged curves are shown for FAK inhibitor-only samples (N = 7), and individual curves are shown for slices with FAK inhibitor and external pressure (N = 4). **(j)** Immunofluorescence of GM130 (GOLGA2) in a chick foregut section. **(k)** Zoomed-in view of the red box in (j) overlaid with mesenchymal cell orientation vectors indicated by colored arrows. 0° marks direction towards the epithelium. **(l,m)** Distributions of cell orientation in control (l, N = 9) and FAK inhibitor-treated (m, N = 7) slice culture. Each colored line represents one biological replicate, and the black line shows the average. The dashed purple line marks the expected uniform distribution of angles for random cell orientation (null hypothesis). Scale bars: 50 µm. Error bars: SD. P values: Chi-square test against uniform distribution.

The above results indicate that the directional flow of mesenchymal cells towards the epithelium may underlie TES. To test the functional relevance of the mesenchymal flow, we inhibited cell migration with the focal adhesion kinase (FAK) inhibitor PF-573228 in the slice culture. FAK is an enzyme essential for cell migration whose activity is particularly high in the mesenchyme (Supplementary Fig. 7d). In both chick and mouse, inhibition of cell migration significantly reduced the mesenchymal flow, in terms of both speed and directionality (Supplementary Fig. 7e,f). In all treated samples, TES was stalled at the epithelial narrowing stage (Fig. 5g-i, Supplementary Fig. 7e,f, Supplementary Movie 16). As with the surgical removal of the mesenchyme, the effect of FAK inhibition can be rescued by external pressure (Fig. 5g-i, Supplementary Fig. 5e, Supplementary Movie 17).

We complemented the live-imaging results with fixed-tissue analysis, using the vector formed by the Golgi apparatus and the nucleus as an indicator of mesenchymal cell polarity^29^ (Fig. 5j,k). We found that the mesenchymal cells collectively oriented towards the epithelium, producing an angular bias in their polarity vectors (Fig. 5l), which was absent in FAK inhibitor-treated samples (Fig. 5m).

In conclusion, through a comprehensive series of biophysical and pharmacological perturbations in the ex vivo slice culture system, we have demonstrated that epithelial splitting in TES is initially driven by the convergent mesenchymal pressure due to directional cell flow, which is critical to narrow the epithelium mediolaterally to form the septum and enable tube splitting.

### SHH signaling is essential for generating the convergent mesenchymal force

A clue to the molecular regulation of the directional cell movements underlying TES came from consideration of congenital birth defects in which the foregut fails to separate and develops as a fused tube. Laryngo-tracheo-esophageal cleft (LTEC) is a relatively rare human anomaly where an abnormal connection between the airway and esophagus exists due to a failure of TES^14,15^. Notably, the LTEC phenotype is closely modeled in mice deficient in SHH signaling, implicating this pathway in achieving normal TES^11,23,30–32^. To investigate this further, we first confirmed that SHH signaling, indicated by the expression of the PTCH1 gene, is widespread in the foregut mesenchyme in both mouse and chick; and that it is lost upon genetic knockout of Shh in the mouse or *in ovo* pharmacological inhibition of SHH signaling in the chick (Fig. 6a, Supplementary Fig. 9a). Notably, the loss of SHH signaling also dramatically reduces the number of mesenchymal cells, with less significant impact on foregut patterning (Fig. 6b, Supplementary Fig. 9b). The mesenchymal hypoplasia is mainly due to a lack of proliferation, rather than excessive cell death (Supplementary Fig. 9c-f). Importantly, mesenchyme removal showed that the mesenchymal compressive force is lost upon SHH inhibition (Fig. 6c-e). Collective cell polarity towards the mesenchyme was also lost with SHH inhibition (Fig. 6f). Live imaging of cyclopamine-treated chick and Shh-knockout mouse foregut slices further revealed that SHH signaling is essential for the narrowing of the epithelium, as its loss leads to stalling or even relaxation of the epithelial morphology (Fig. 6g-i, Supplementary Fig. 9g, Supplementary Movie 18). Application of external pressure could partly rescue the LTEC phenotype, allowing the formation of the epithelial septum between the forming esophagus and airway, although the resolution of the septum took significantly longer than in wildtype, implying a role for SHH signaling in epithelial remodeling as well (Fig. 6g, Supplementary Movie 19). PIV analysis further showed that SHH signaling is necessary for the convergent flow pattern of the mesenchyme (Supplementary Fig. 9h).

**Figure 6.**
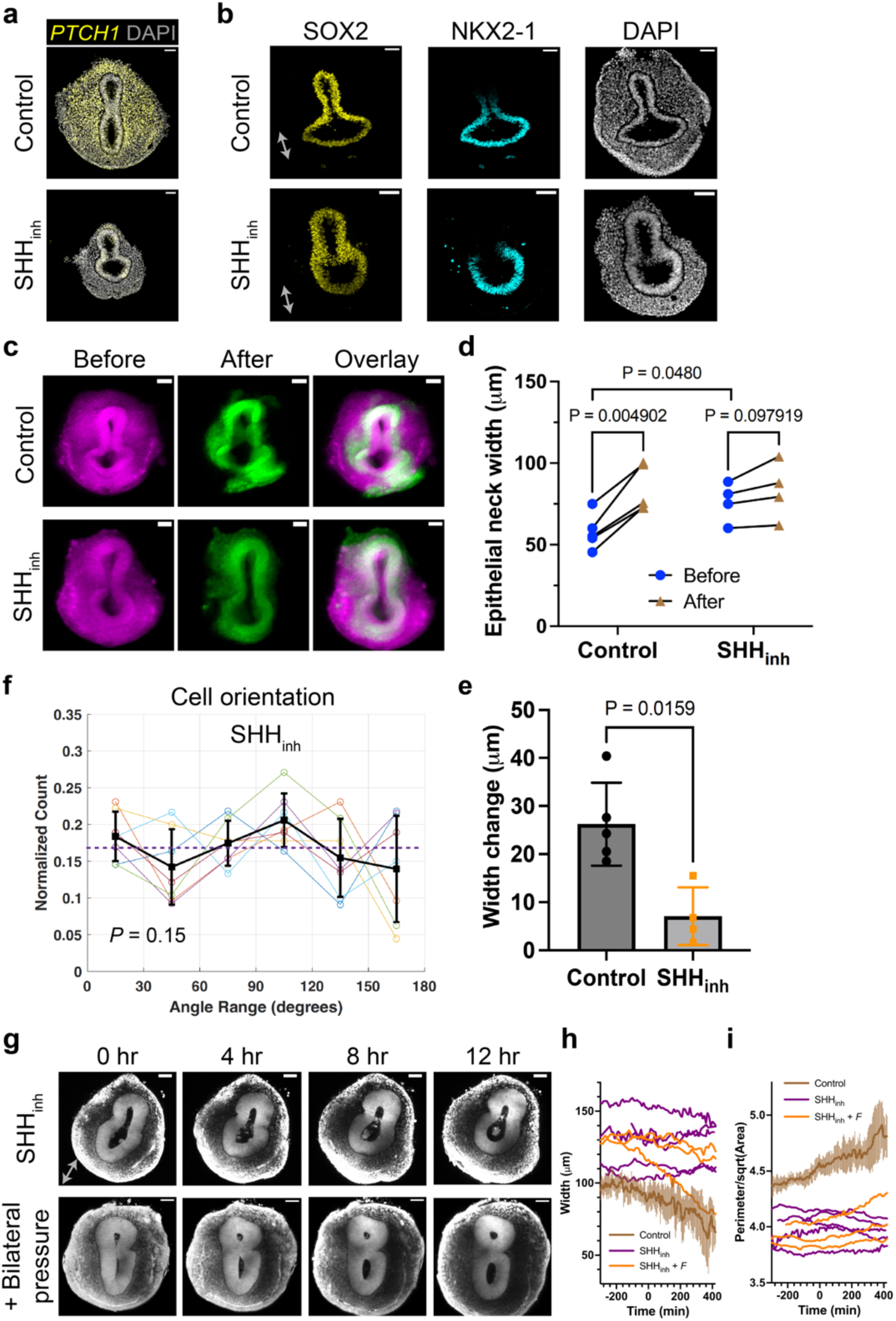
SHH signaling is essential for generating the convergent mesenchymal force. **(a)** HCR-FISH of PTCH1 in transverse sections of E4.0 chick foreguts with or without *in ovo* SHH inhibitor (cyclopamine) treatment. **(b)** Immunofluorescence of SOX2 and NKX2-1 in transverse sections of E3.75 chick slices with or without *in ovo* SHH inhibitor treatment. **(c)** Stereoscope imaging of a control (upper) and a SHH inhibitor-treated (lower) GFP chick foregut slice before (magenta) and after (green) surgical removal of the mesenchyme. **(d)** Quantification of epithelial neck widths (defined in Fig. 3a) before and after mesenchyme removal (N = 4 or 5 samples). **(e)** Comparison of epithelial neck width changes calculated from (d). **(f)** Distributions of mesenchymal cell orientation in SHH inhibitor-treated slice culture as in Fig. 5l (N = 7 samples). **(g)** Live imaging of E3.75 GFP chick slices treated with SHH inhibitor *in ovo*, in the absence (top) or the presence (bottom) of bilateral pressure by a collagen gel trough. **(h,i)** Quantification of the neck width (h) and the perimeter-area ratio (i) of SHH inhibitor-treated slices without (purple) or with external bilateral force (orange) compared with E3.75 controls (brown, as in Fig. 3c,e). Each curve shows an individual sample (N = 5 for SHH inhibitor-only samples, and N = 3 for SHH inhibitor with bilateral pressure). Scale bars: 50 µm. Error bars: SD. P values: Multiple paired t test (d), Mann-Whitney test (e), Chi-square test against uniform distribution (f).

### The dorsal sub-epithelial mesenchyme is sensitive to SHH signaling and contributes to the mesenchymal force

Recent studies uncovered multiple cell types in the developing foregut mesenchyme which could be potentially responsible for driving the SHH-dependent convergent flow^23,33^. To understand which cell type responds to SHH signaling to enact the directed movement, we performed single-cell RNA sequencing (scRNA-seq) of foregut slices from normal and cyclopamine-treated chick embryos and used graph-based clustering to establish cell groupings putatively representing distinct cell types (Fig. 7a). We then spatially mapped the identified cell populations in the foregut by staining for marker genes representative of each cluster (Fig. 7b, Supplementary Fig. 10a,b,11a). We found that the foregut mesenchyme is composed of sub-epithelial and peripheral populations along the radial axis from the epithelium, in addition to the dorsoventral axis corresponding to the esophageal and tracheal cell types. Upon SHH inhibition, the cell population whose size is most affected is the dorsal sub-epithelial mesenchyme, whereas the proportions of other major cell populations remain balanced (Fig. 7c,d, Supplementary Fig. 10c). The dorsal sub-epithelial mesenchyme, marked by NKX6-1, also express SHH response genes including PTCH1/2 and FOXF1/2, which are dramatically downregulated with cyclopamine treatment (Fig. 7e). Although smooth muscle differentiation has not yet begun during TES, we found early expression of smooth muscle actin (ACTA2) in a subset of dorsal sub-epithelial mesenchyme, which plummeted upon SHH inhibition (Supplementary Fig. 10d). This may underlie the smooth muscle defects observed in SHH mutants^34,35^. Gene ontology analysis of differentially expressed genes between control and cyclopamine-treated dorsal sub-epithelial mesenchyme revealed that pathways related to cell migration, including extracellular matrix (ECM)-receptor interaction, axon guidance, and focal adhesion, are significantly downregulated with SHH inhibition, suggesting a role of SHH in directing cell migration (Fig. 7f). As SHH is secreted by the epithelium and sensed by the mesenchyme, we further analyzed intercellular communication pathways between dorsal epithelium and dorsal sub-epithelial mesenchyme/peripheral mesenchyme (Supplementary Fig. 11b). Among the predominant signaling pathways between these cell types, including SHH, semaphorin, and specific ECM-receptor interactions, only SHH signaling is dramatically weakened by cyclopamine treatment (Supplementary Fig. 11b). Immunostaining confirmed that the dorsal sub-epithelial mesenchyme is almost absent with *in ovo* cyclopamine treatment, with the peripheral mesenchyme marked by TBX1 surrounding the dorsal epithelium (Fig. 7g). In such case, the remaining dorsal peripheral mesenchyme is insufficient to generate the compressive force critical to TES.

**Figure 7.**
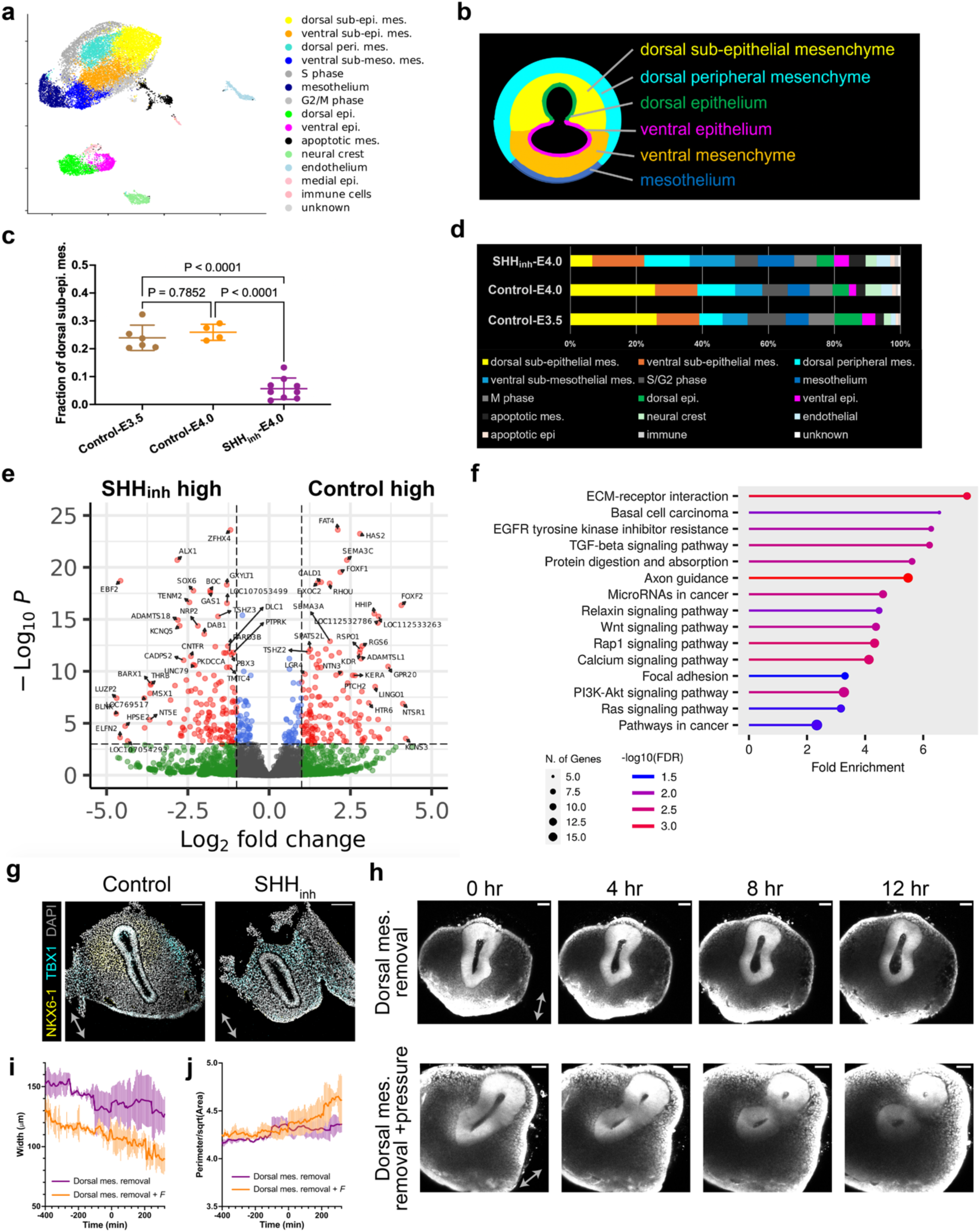
Single-cell RNA sequencing identifies the SHH-responsive dorsal sub-epithelial mesenchyme as the driver for TES. **(a)** UMAP of single-cell transcriptomes in E3.5 and E4.0 chick foregut slices. Numbers indicate different cell clusters. **(b)** Schematics of the spatial distribution of major epithelial and mesenchymal cell types in the foregut based on HCR-FISH and immunofluorescence of select marker genes. **(c)** Fractions of the dorsal sub-epithelial mesenchyme in individual embryos recovered from scRNA-seq data. Each data point represents one embryo. P values are calculated by one-way ANOVA with Dunnett’s test. **(d)** Cell composition analysis of E3.5 control, E4.0 control, and E4.0 cyclopamine-treated samples. **(e)** Volcano plot highlighting differentially expressed genes (DEG) between control and cyclopamine-treated samples. Genes with log2-fold change > 1 and *P_adj_* < 10^-4^ are marked red. **(f)** Gene ontology analysis of downregulated DEGs by SHH inhibition. **(g)** Immunofluorescence of NKX6-1 and TBX1 in transverse sections of E4.0 chick foreguts with or without *in ovo* cyclopamine treatment. **(h)** Live imaging of an E3.75 GFP chick slice with surgical removal of the dorsal mesenchyme in the absence (upper) or presence (lower) of bilateral pressure. **(i,j)** Quantification of the neck width (i) and the perimeter-area ratio (j) of dorsal mesenchyme-removed slices without (purple) or with external bilateral force (orange). N = 6 or 8 samples. Scale bars: 50 µm. Error bars: SD.

These results suggest that the SHH is essential for the specification and proliferation of the dorsal sub-epithelial mesenchyme, the loss of which leads to defective mesenchymal force, and hence a failure to enable TES. To directly test the functional importance of the dorsal mesenchyme, we surgically removed the dorsal mesenchyme while keeping the ventral part intact (Fig. 7h, Supplementary Movie 20). Strikingly, the removal of only the dorsal mesenchyme was sufficient to halt TES, verifying that the major source of the compressive mesenchymal force is the dorsal mesenchyme (Fig. 7h). Quantitative analysis shows that although the ventral mesenchyme was still present, the initial epithelial neck width increased and its reduction slowed (Fig. 7i, Fig. 3c), both of which were rescued by applying additional bilateral pressure (Fig. 7j, Supplementary Movie 21). These results suggest a significant and essential contribution of mesenchymal force from the dorsal mesenchyme. The presence of medial constriction rather than dorsal compression can be explained by the dorsal mesenchymal domain extending beyond the prospective septation point along the dorsoventral axis and by the substantially greater number of cells surrounding the epithelium bilaterally than dorsally (Fig. 7g and Supplementary Fig. 10b).

### SHH induces directional migration of the foregut mesenchymal cells to deform the epithelium

Our results showed that SHH is essential for the specification and proliferation of the dorsal sub-epithelial mesenchyme, which in itself could potentially generate static crowding pressure on the epithelium. To test whether, in addition, SHH plays a role in inducing directional mesenchymal cell migration essential to TES, we implanted beads loaded with SHH ligand into the foregut mesenchyme (Fig. 8a). Subsequently, 24 hours post-implantation, the mesenchyme near the SHH bead showed significantly higher density than the contralateral side, whereas implantation of an empty bead had no such effect (Fig. 8b). The increased cell density is not due to enhanced cell proliferation by SHH because the fraction of dividing cells was not increased near the beads (Supplementary Fig. 12a,b), potentially because the endogenous SHH level was sufficient to sustain a relatively high proliferation rate. Live imaging further revealed that mesenchymal cells near the bead moved toward the SHH source, instead of moving towards the epithelium on the contralateral side (Fig. 8c-e, Supplementary Movie 22). Such effect was not reproduced with an empty bead (Supplementary Fig. 12c-f, Supplementary Movie 23). The cell polarity vectors showed a significant bias towards the SHH bead at an ∼60° angle (Fig. 8f-h), potentially due to cell packing density, which prevented unlimited persistent migration toward the bead. These data strongly suggest that foregut mesenchymal cells are attracted to the source of SHH activity.

**Figure 8.**
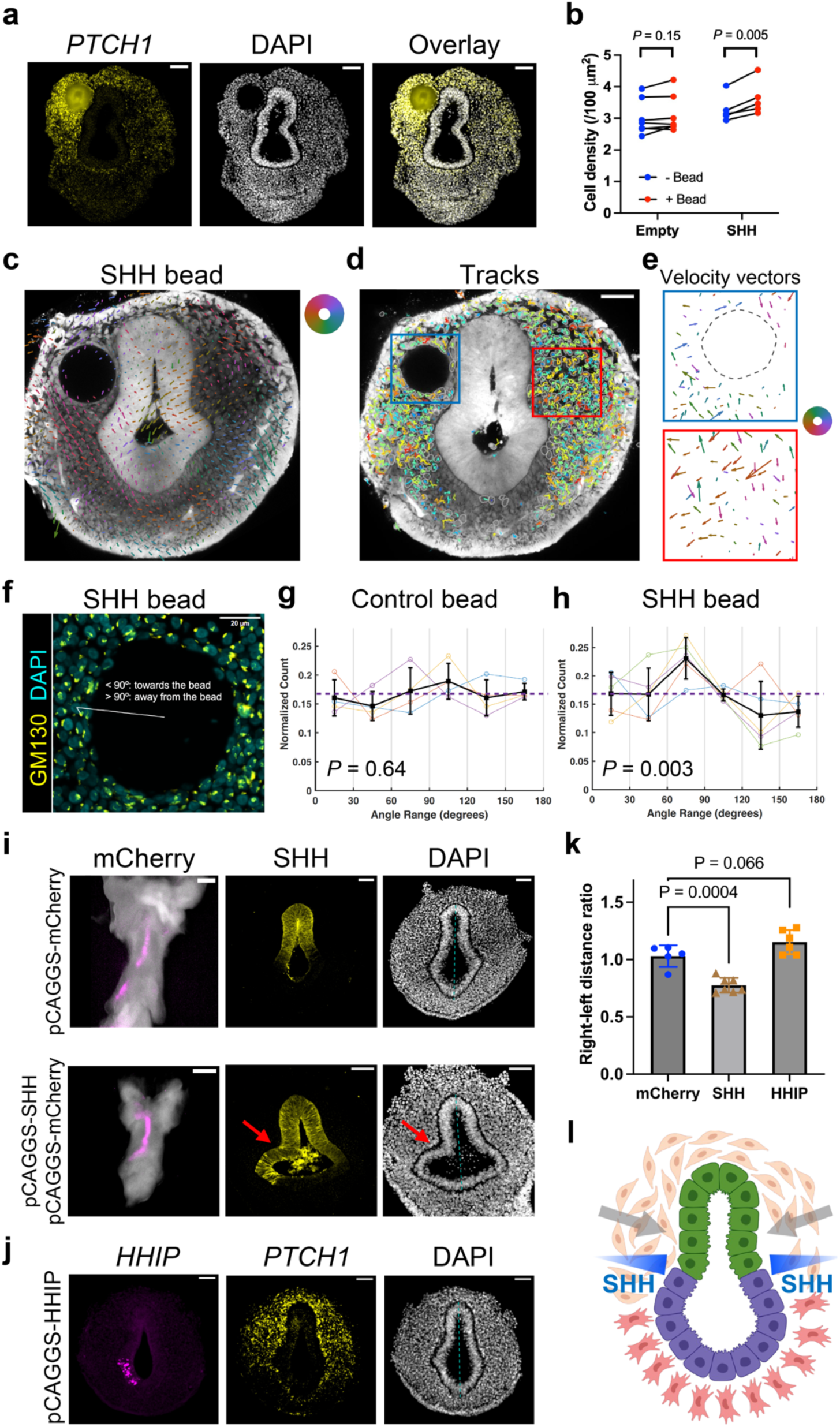
SHH attracts the foregut mesenchymal cells to deform the epithelium. **(a)** HCR-FISH of *PTCH1* in an E3.5 chick foregut slice 24 hours after implantation of an Affi-Gel bead loaded with recombinant mouse SHH protein. **(b)** Comparison of mesenchymal cell density near the implanted bead (with or without SHH) and the contralateral side. Each dot represents one slice sample. **(c)** PIV analysis of an E3.75 GFP chick slice with an implanted SHH-bead. The color wheel indicates the directions of PIV vectors. **(d)** Live imaging of a SHH-bead-implanted GFP chick slice overlaid with cellular trajectories colored by their average speeds. **(e)** Average velocity vectors of individual cell trajectories in the blue and red boxes of (d). Vectors are scaled with the speed and color-coded by their directions according to the color wheel. The dashed circle marks the bead. **(f)** Immunofluorescence of GM130 (GOLGA2) in a chick foregut section with a SHH-loaded bead. **(g,h)** Distributions of mesenchymal cell orientation in slices implanted with control beads (g, N = 4) and SHH beads (h, N = 6). **(i,j)** *In ovo* electroporation of plasmids encoding mCherry (i), SHH (i), or HHIP (j) into the right epithelium of the chick foregut. Frontal views of the foreguts by stereo microscopy shows the effective electroporation (i). Immunofluorescence of SHH (i) and HCR-FISH of *HHIP* and *PTCH1* (j) confirm the efficient expression of the constructs. Arrows indicate the asymmetrical deformation of the epithelium induced by SHH overexpression. The dashed lines bisect the foregut lumen as a reference for measuring epithelial distances to the midline. **(k)** Comparisons of right (electroporated) and left (non-electroporated) epithelium-to-midline distances with control (mCherry), SHH, and HHIP electroporation. **(l)** Working model of TES initiation. SHH secreted from the foregut epithelium induces dorsal sub-epithelial mesenchyme’s migration towards the epithelium, generating bilateral compressive force essential to TES. Scale bars: 50 µm (except the left panels in (i) which are 100 µm). Error bars: SD. P values: Paired t test (b), Chi-square test against uniform distribution (g,h), One-way ANOVA with Dunnett’s test (k).

Next, we attempted to perturb the symmetry of SHH expression to test whether the strength of SHH signaling correlates with epithelial deformation. When we overexpressed SHH, but not mCherry as negative control, in the right foregut epithelium by electroporation, the right foregut epithelium was more deformed than the left side, implying the induction of a stronger mesenchymal force with the increase of SHH signaling (Fig. 8i). Conversely, when the SHH antagonist Hedgehog Interacting Protein (HHIP) was overexpressed, SHH signaling near the electroporated site was attenuated, resulting in a slightly less deformed epithelium compared to the unaffected side, potentially due to the inability of HHIP to fully inhibit SHH secretion (Fig. 8j,k). Therefore, induction of directional cell migration by physiological SHH signaling is sufficient to initiate epithelial deformation for tube splitting. It is possible that the mesenchymal cells are chemotactic to SHH secreted by the epithelium, and as the epithelium constricts, a negative curvature then concentrates SHH to generate positive chemo-mechanical feedback which facilitates epithelial splitting.

## Discussion

In search for the biomechanical basis of epithelial tube splitting, we focused on TES, a critical epithelial splitting event in foregut development. We established an ex vivo slice culture system to observe and perturb TES with single-cell resolution. This enabled us to uncover an unappreciated role of mesenchymal force due to convergent cell migration. Gaining insights from mouse models and human patients with defective TES, we identified SHH signaling as both the driver of the convergent cell flow and the maintainer of mesenchymal proliferation, which is essential for TES. Our results provide a biomechanical mechanism bridging the gap between the existing genetic understanding of TES and its morphogenetic outcome, demonstrating how SHH signaling from the epithelium to the mesenchyme transforms into convergent mesenchymal force driving TES (Fig. 8l).

### Epithelial tube splitting morphogenesis requires mesenchymal force

The mechanism of epithelial folding has attracted the attention of biophysicists and developmental biologists for decades. Early work highlighted that the asymmetric distribution of actomyosin within the epithelium can drive folding such as in neural tube closure and intestinal crypt morphogenesis^36,37^. Anisotropic proliferation of the epithelial cells is also known to fold the epithelium through buckling^27,38^. Besides the epithelium-intrinsic mechanisms, we have just begun to appreciate the biomechanical role of the adjacent mesenchyme, whose signaling role in epithelial-mesenchymal crosstalk is much better understood^5,6,39–41^. For example, spontaneous clustering of mesenchymal cells underlies the initial intestinal villus formation in the mouse^7,42,43^. Although either epithelium-intrinsic or -extrinsic forces can fold the epithelium in various systems, it remains unknown whether either, or both forces, is sufficient to drive epithelial tube splitting, which starts with high-curvature folding and ends up with a topological transition in the epithelium. The poor understanding of epithelial splitting is at least in part due to the lack of a tractable experimental system. TES, where the dorsoventrally patterned foregut epithelium splits into the dorsal esophagus and the ventral trachea, is among the few biological examples of epithelial splitting^8,10^. The esophageal and tracheal progenitors sort into distinct domains via actomyosin contractility downstream of heterotypic Ephrin signaling, yet in vitro reconstituted cell domains remain connected, suggesting additional force is required for initiating epithelial splitting^13,44^.

Our ex vivo slice culture system recapitulates the tissue dynamics of TES characterized in vivo, and importantly, allows cross-sectional imaging with single-cell resolution. By surgical and pharmacological manipulations of the foregut mesenchyme in slice culture, we demonstrated that the cell flow-dependent mesenchymal force is crucial to epithelial narrowing in TES to form the epithelial septum. The septal epithelial cells then undergo apicobasal remodeling and apoptosis to resolve into two tubes^11,24^. It is noteworthy that the critical role of mesenchymal force does not imply that epithelium-intrinsic processes are unimportant. Indeed, previous genetic studies have identified essential epithelial mechanisms in TES, especially the proper patterning of the SOX2 and NKX2.1 domains^13,20,45^. Therefore, TES requires both mesenchymal and epithelial forces, which may also hold true in other epithelial splitting systems, such as cloacal septation of the embryonic hindgut.

### SHH signaling integrates molecular patterning of the foregut and TES morphogenesis

Beyond elucidating the biomechanical basis of TES, we also sought to connect the biophysical mechanism to the existing knowledge about foregut patterning through genetic studies. It was demonstrated almost two decades ago that the gradients of BMP and Wnt signaling in the mesenchyme instruct the dorsoventral patterning of the foregut epithelium into SOX2^+^ esophageal and NKX2-1^+^ ventral domains, which secretes SHH to sustain the mesenchyme^19,22,23,46,47^. Deficient SHH signaling has been observed in human patients with TES defects as well as in genetic mouse and frog models, where the foregut mesenchyme exhibits various degrees of hypoplasia depending on the severity of SHH reduction^11,48^. Although these studies have implicated a role for the mesenchyme in TES morphogenesis, the mechanisms have not been fully investigated. In our mouse and chick models of SHH deficiency, the foregut epithelium fails to narrow bilaterally to initiate TES, in a similar fashion to the mesenchyme-reduced wildtype sample in slice culture. Combining scRNA-seq, bead implantation, and targeted electroporation, we showed that the dorsal sub-epithelial mesenchyme is the main effector of SHH in TES morphogenesis. SHH signaling can directly induce mesenchymal cell migration and is crucial for dorsal mesenchymal cell proliferation^49–51^. Both roles are essential to TES. Inhibition of the migratory force by the FAK inhibitor fully blocks TES (Fig. 5g-i), whereas a reduction in mesenchymal cell number by surgical dissection can also delay TES (Fig. 7h-j). However, blocking cell proliferation at later stages does not prevent TES (Fig. 4), suggesting a critical threshold of mesenchymal cell number in TES. SHH signaling in the mesenchyme leads to a convergent bilateral force towards the epithelium, creating the septum at the dorsoventral boundary of the epithelium which is most mechanically susceptible, potentially due to ephrin-mediated cell segregation. Our results thus bridge the gap between molecular patterning and tissue morphogenesis, demonstrating how epithelial-mesenchymal interactions function synergistically to enable robust epithelial splitting.

### Mechanobiology of morphogenesis sheds light on the etiology of structural birth defects

In the study of structural birth defects, much of the focus has been on evaluating individual disease-causing genes, and their roles in normal morphogenesis; taking advantage of the rapidly expanding base of clinical genetics data as a starting point^52,53^. While this paradigm has provided valuable insights into the molecular underpinnings of many conditions, critical aspects of the underlying mechanisms might be missed when pertinent genes have pleiotropic effects essential for embryonic development or early postnatal life. For example, although genetic disruptions of SHH signaling (Shh, Gli2), WNT signaling (Wnt2/2b), and Ephrin signaling (Efnb2) cause TES defects in mouse embryos with nearly 100% penetrance, these pathways are underrepresented or absent in the clinical data of TES-related birth defects, likely because these pathways are critical for other aspects of embryonic development^14,15^. Consequently, studying normal tissue morphogenesis is crucial for understanding the full spectrum of factors contributing to proper organ formation. Mechanobiological processes, in particular, play an essential role in shaping tissues and organs during development, but these mechanisms remain underappreciated in the development of many organ systems. By focusing on the mechanobiology of physiological and pathological TES morphogenesis, our work unveils the vital role of mesenchymal force in shaping the foregut epithelium, broadening our understanding of the etiology of foregut birth defects.

## Acknowledgements

We thank the MicRoN Facility of Harvard Medical School for offering equipment and training for microscopy experiments, the Biopolymers Facility of Harvard Medical School for performing next-generation sequencing, and the Center for Comparative Medicine of Harvard Medical School for maintenance of the mouse colonies. We thank members of the Tabin group and the Mahadevan group for discussion and suggestions. This work was funded by the NIH grant HD087234 (C.J.T.), the Helen Hay Whitney Fellowship (R.Y.), the NWO Rubicon grant (L.A.H.), the Simons Foundation (L.M.), and the Henri Seydoux Fund (L.M.).

## Author contributions

R.Y. conceived the project. R.Y. and C.J.T. designed the experiments and wrote the draft of the manuscript. R.Y. performed all experiments, analyzed the data, and wrote the codes for plots figures. L.A.H. performed analysis and quantification of the foregut morphology in slice culture and gave advice on image processing and PIV analysis. P.O. wrote the code for computing the strain rate tensor from the PIV data. D.L. wrote the code for SAM 2 segmentation. C.L. performed initial processing and analysis of the single-cell RNA-sequencing data. H.K.G. gave advice and technical support on *in ovo* electroporation and slice culture. A.M., N.L.N., and L.M. gave advice on experimental design and data analysis. L.M. and C.J.T. supervised the study. All authors read the manuscript and gave suggestions.

## Methods

All animal studies were performed in compliance with the protocols approved by the Institutional Animal Care and Use Committee at Harvard Medical School.

### Chick embryos

Fertilized Specific Pathogen Free (SPF) white Leghorn chicken eggs (Charles River/AVS Bio) and transgenic Roslin Green (expressing cytoplasmic GFP) chicken eggs (Susan Chapman, Clemson University) were used. Eggs were incubated in a 38°C humidified chamber before collection.

### Mouse embryos

All mouse lines used were obtained from the Jackson Laboratory: C57BL/6J (#000664), *ROSA^nT-nG^* (*nTnG*, #023537), *Shh-GFP-Cre* (#005622), and *Shh-CreERT2* (#005623). We crossed *Shh-CreERT2* with *nTnG* to generate the strain *ShhCreER/+;nTnG/nTnG*, which was then crossed with *Shh-GFP-Cre* to generate the *Shh*-knockout embryos and littermate controls. All mice were housed in a specific pathogen-free (SPF) facility at Harvard Medical School under a 12-hour light/12-hour dark cycle. Animals had ad libitum access to food and water. Timed pregnancy was set up for all experiments. The morning when a vaginal plug was observed was designated as E0.5.

### Plasmids

pCAGGS-mCherry was a gift from Phil Sharp (Addgene plasmid #41583). pCAGGS-SHH and pCAGGS-HHIP was made by replacing mCherry with chick SHH (cloned from Addgene plasmid #13991) or mouse HHIP (Origene, MC203592) using restriction/ligation cloning. pCAGGS-MLC2-tdTomato was made by cloning mouse MLC2-tdTomato (from Addgene plasmid #58108) to pCAGGS. The plasmids will be deposited to Addgene. QIAGEN Plasmid Maxi Kit was used to purify plasmids from in-house prepared DH5-alpha cells.

### Foregut slice culture

Fresh chick or mouse embryos were collected and promptly dissected in chilled Dissection Medium (4% fetal bovine serum [Peak Serum, PS-FB2] and 100X-diluted penicillin-streptomycin [Gibco, 15140-122] in DMEM with HEPES [Gibco, 21063029]) to isolate the foregut, from the pharyngeal arches to the upper stomach. The foregut was then manually sliced with a surgical blade (Aspen Surgical, 371111) transversally to obtain slices of 100-150 µm thickness. The slice containing the septum and the anterior undivided foregut was embedded in 50 µL 50% Matrigel (Corning, 356231) diluted with the Dissection Medium on a 35-mm glass bottom dish (MatTek, P35G-1.5-14-C), with the undivided foregut side facing up. Once the slice was settled in Matrigel at room temperature, the dish was incubated at 37°C in a humidified chamber for 30 minutes to solidify the Matrigel. 2-mL of culture medium for chick (5% chick embryo extract [US Biological, C3999], 3% fetal bovine serum, 0.25 µM LDN-193189 [Cayman Chemical, 11802], and 100X-diluted penicillin-streptomycin in FluoroBrite DMEM [Gibco, A1896701]) or mouse (100 ng/mL EGF [R&D Systems, 236-EG], 5% fetal bovine serum, 0.25 µM LDN-193189, and 100X-diluted penicillin-streptomycin in FluoroBrite DMEM) was carefully added to the dish, and the sample was incubated at 37°C with 5% CO_2_ prior to imaging.

### Surgical removal of foregut mesenchyme

Foregut slices were trimmed with fine forceps (Fine Science Tools, 11252-00) to reduce the mesenchyme without impacting the epithelium. Stereo images before or immediately after mesenchyme removal were taken on a Leica stereoscope (M165 FC) with an LED light source (Lumencor, SOLA), and the images were aligned using a custom MATLAB code. The removal of chick foregut mesenchyme can be nearly complete as the tissue is relatively large and the epithelium is robust, whereas for the mouse foregut we usually leave some mesenchyme to avoid tearing and breaking the fragile epithelium.

### Laser ablation

Laser ablation was performed with the Leica Stellaris 8 multiphoton microscope with the Insight X3 dual beam laser and the 25X/1.0NA water immersion objective. GFP chick slices were excited at 960-nm with 7% power and FLIPPER-TR-labeled mouse slices were excited at 1040-nm with 7% power. A video before ablation was first recorded as control. Laser ablation was then performed with 800-nm excitation at 70%-80% power depending on the depth of the imaging plane, in a defined ROI marking the sub-epithelial mesenchyme. The ROI was ablated for ∼0.1 seconds per 4-second frame for 3-10 frames, with transmission light image being recorded during ablation to monitor the damage of cells. Once the cells were ablated, the system was switched to the imaging mode as before ablation with a recording rate of 4-12 seconds per frame.

The post-ablation video was drift-corrected with the Fiji StackReg plugin (https://bigwww.epfl.ch/thevenaz/stackreg)^54^. Kymographs were generated with the Multi Kymograph function in Fiji (https://biii.eu/multi-kymograph) along lines normal to the contour of the epithelium crossing the ablated region.

We quantified three characteristics of recoil after ablation. First, the initial displacement rate was determined by the slope of the tangent line at the kymograph’s base, which reflects the force imbalance at the tissue interface and the epithelial viscosity. Second, we measured the total epithelial displacement by tracking the movement of the basal side of the epithelium, which is a consequence of force imbalance and epithelial elasticity^55^. However, quantitative fitting of the tissue displacement by a simple Kevin-Voigt model is complicated by the simultaneous expansion of the epithelial width in both directions due to the lack of resisting force on the apical side of the epithelium.

### Time-lapse imaging of the foregut slice culture

GFP chick slices were excited at 960-nm with 7% power and nTnG mouse slices were excited at 1040-nm with 7% power on the abovementioned two-photon microscope. A Z stack of 30-60 µm with 10-15 µm step size was recorded for each slice, starting from ∼30 µm below the tissue surface to avoid potential artifacts from cutting. Samples were imaged every 10-30 minutes for 12-16 hours. The XY resolution ranged from 250 nm to 500 nm per pixel. The time-lapse videos were used for quantitative analysis of the morphological evolution of the foregut (see “Morphometric analysis of the foregut epithelium”).

### Bilateral compression of foregut slices

We made a narrow trough in a collagen gel pad to compress the foregut slice. The collagen gel was made with a solution containing 2 mg/mL rat tail collagen (Gibco, A1048301), 1X PBS (Invitrogen, AM9625), and 17 µM sodium hydroxide (Supelco, SX0607N) in water. 1 mL collagen solution was spread on a 35-mm glass bottom dish and incubated at 37°C for 30 minutes. The solidified gel pad was buffered at room temperature with 2 mL Dissection Medium, then 1 mL culture medium. A 1-mm long, 200-300-µm wide trough was made in the gel pad with a fine tungsten needle (Fine Science Tools, 10130-05), and the foregut slice was gently squeezed bilaterally with forceps to fit into the trough with the dorsoventral axis parallel to the long axis of the trough. The sample was incubated at 37°C for several hours before live imaging as described.

We also used the collagen gel to calibrate the stability of the imaging setup. We embedded 200-nm red fluorescent beads (Invitrogen, F8810) at 1:10,000 dilution in collagen solution prior to gelation. During live imaging, the beads were imaged with the 1040-nm laser and tracked with TrackMate to generate the trajectories.

### Pharmacological treatment of foregut slices

We used drugs at concentrations that effectively inhibited the biological target with minimal adverse effects on the tissue integrity and viability. 3 µM aphidicolin (Tocris, 5736) or 2.5 µM PF-573228 (MedChemExpress, HY-10461) was added to the slice culture medium 2-3 hours prior to live imaging for 12-24 hours.

### *In ovo* cyclopamine treatment

To inhibit SHH signaling in vivo, we developed an *in ovo* cyclopamine treatment protocol for chick embryos. 3 mL of albumen was drawn from the bottom of the egg at E2 to prevent the embryo from sticking to the eggshell. The egg was sealed with tape and incubated to E3 when it was windowed. 50 µL 10 mM cyclopamine (ApexBio, A8340) dissolved in ethanol was mixed with 250 µL PBS by gentle pipetting, and the mixture, usually in a form of cloudy suspension, was carefully added on top of the chick embryo. The treated egg was immediately sealed and returned to the incubator. We found that this protocol effectively inhibited SHH signaling, yielding phenotypes highly similar to the *Shh*-knockout mouse embryos. The timing and dose of cyclopamine addition could be tuned to inhibit SHH signaling in a controlled manner.

### Bead implantation of foregut slices

A 20-µL aliquot of 100-200 mesh Affi-Gel blue beads (Bio-Rad, 1537302) was rinsed with 1.5 mL PBS overnight at 4°C. 10 µL of 2 mg/mL recombinant mouse SHH N-terminus (R&D Systems, 464-SH) was dropped on the lid of a plastic dish placed on ice. Due to the small size of the foregut tissue, we picked the smallest beads to transfer into the SHH droplet, incubating on ice for 1-2 hours. The loaded beads were implanted to the dorsal mesenchyme of foregut slices using fine forceps, making sure the implanted bead was in the middle of the slice to prevent dislodging of the bead. Implanted slices were then embedded and cultured in Matrigel as describe above. As control, the PBS-rinsed beads were transferred to a droplet of PBS on ice prior to implantation.

Slices were fixed 24 hours post-implantation for HCR of chick PTCH1. To quantify mesenchymal cell density, we counted the number of DAPI-positive nuclei within a ring 10 µm to 50 µm away from the bead surface, and then divided the total cell number by the area of the ring. On the contralateral side without the bead, a square of equivalent area was drawn to count the cells.

### *In ovo* electroporation of the foregut epithelium

The plasmid mixture for electroporation was made with 4 µg/µL plasmid, 0.5% Fast Green FCF (Sigma-Aldrich, F7252), and 3% sucrose (Sigma-Aldrich, S8501) in TE buffer. E2 chick embryos (HH10-12) were lowered and windowed. The embryo was counterstained with an injection of ∼30 µL 20X diluted ink (Pelikan, 211862) in PBS to the yolk below the embryo. A mouth pipette with a thin tip, made with glass capillary (FHC Inc, 30-30-0) pulled by a micropipette puller (Sutter Instrument, P-97), was loaded with the plasmid mixture, which was injected into the foregut lumen dorsally until the dye flowed out from the anterior intestinal portal. A parallel needle electrode (Bulldog Bio, CUY560-5-0.5) was inserted into the yolk parallel to the anterior-posterior axis of the embryo, with the positive electrode on the right side of the embryo. Electroporation was performed with three 50-V, 8-ms, 100-ms apart poring pulses, followed by five 20-V, 8-ms, 100-ms apart transfer pulses, all towards the positive electrode (Nepa Gene, NEPA21). A thin layer of fresh albumen was added to the top of the embryo to prevent drying, and the egg was taped and incubated to the desired stage.

### EdU labeling and detection

10 µM EdU (5-ethynyl-2-deoxyuridine, Invitrogen, A10044) was added to the culture medium for foregut slices. After two hours of incubation at 37°C, the sample was fixed, embedded, and sectioned as described in “Immunofluorescence of tissue sections”. EdU labeling was performed with the Click-iT EdU Cell Proliferation Kit for Imaging with Alexa Fluor 488 (Invitrogen, C10337) according to the manufacturer’s protocol.

To quantify the fraction of EdU-positive cells, we manually counted the total number of cells labeled by DAPI and the EdU-positive cells in 30X30 µm^2^ boxes within the dorsal, medial, or ventral mesenchyme. At least three boxes were counted for each region in each slice, and the results were averaged as one biological replicate.

### Whole-mount immunofluorescence

Whole embryos (chick embryos before E3.5 and mouse embryos before E10.0) or dissected foreguts (chick embryos after E4.0 and mouse embryos after E10.5) were fixed in 4% paraformaldehyde (PFA, Electron Microscopy Sciences, 15710) diluted in PBS (Gibco, 10010023) overnight at 4°C. The samples were then washed three times with PBS (10 minutes each at room temperature), and blocked with Blocking Buffer (10% bovine calf serum [Gibco, 16010159] and 1% Triton X-100 [Sigma-Aldrich, T8787] in PBS) for one hour at room temperature. Mouse anti-E-Cadherin (BD Biosciences, 610182) was diluted 1:100 in the Blocking Buffer and labeled the samples for two days at 4°C. The samples were then washed three times with the Blocking Buffer (1 hour each at room temperature) and stained with donkey anti-mouse-Cy3 (Jackson ImmunoResearch, 715-165-150) diluted 1:400 in the Blocking Buffer for one day at 4°C. After six times of washing with 5X diluted Blocking Buffer, the samples were serially dehydrated with 50% methanol (Fisher Scientific, A433P-4)/50% PBS, 80% methanol/20% water (Invitrogen, 10977015), and 100% methanol (1 hour each step at 4°C). Optical clearing was performed with the CytoVista Tissue Clearing kit (Invitrogen, V11322) per the manufacturer’s instruction.

Cleared samples were transferred to a 50-mm glass bottom dish (MatTek, P50G-1.5-30-F) filled with the CytoVista Tissue Clearing Enhancer solution (part of the Tissue Clearing kit) and imaged on a Nikon Ti inverted microscope with a W1 spinning disk scanner (Yokogawa, CSU-W1) using a 20x objective (Nikon, MRD70270) and 561-nm excitation. A Z-stack of ∼0.6-1.0 µm step size was acquired for each sample. Three-dimensional rendering of the image was performed in the Arivis Vision4D software (Zeiss), with the Z step size multiplied by 1.5 to correct for the refractive index mismatch. Markers were manually set at the pharyngeal arch-foregut junction, tracheal-esophageal septum, and the tracheal-bronchial junction to measure the undivided foregut length and the trachea length.

### Immunofluorescence of tissue sections

Whole embryos, dissected foreguts, or ex vivo cultured foregut slices were fixed in 4% paraformaldehyde diluted in PBS overnight at 4°C. Fixed tissue was washed three times with PBS (10 minutes each at room temperature), and cryopreserved with a series of 15% sucrose (Sigma-Aldrich, S8501) in PBS, 30% sucrose in PBS, and 1:1 O.C.T. Compound (Sakura Finetek, 4583):30% sucrose in PBS (1 hour each at room temperature). Samples were transferred to and embedded in O.C.T. Compound, frozen in the dry ice-ethanol bath before stored at -80°C. Frozen samples were transversally sectioned using a cryostat (Leica, CM3050 S) to 14-20 µm thickness onto Superfrost Plus glass slides (Fisher Scientific, 12-550-15). The sample slides were baked at 50°C for 20 minutes to dry, and then rinsed three times with PBS (5 minutes each at room temperature). For labeling phospho-MLC, TBX1, and SHH, antigen retrieval was performed with 1X citrate buffer (Abcam, 64214) in a boiling steamer (Amazon, B00DPX8UBA) for 10 minutes. The sample was then cooled down to room temperature and rinsed two times with PBS (5 minutes each at room temperature). For immunolabeling, the samples were first permeabilized with 0.1% Triton X-100 in PBS for 20 minutes, and blocked with the Blocking Buffer (5% donkey serum [Jackson ImmunoResearch, 017-000-121] and 0.3% Triton X-100 in PBS) for one hour. The primary antibodies were diluted in the Blocking Buffer and incubated the sample overnight at 4°C. The labeled samples were washed with the Washing Buffer (10X-diluted Blocking Buffer in PBS) three times (10 minutes each at room temperature), and labeled with secondary antibodies (including 10 µg/mL DAPI [Invitrogen, D1306] or 1:100 phalloidin-Alexa Fluor 488 [Invitrogen, A12379]) diluted in the Blocking Buffer and incubated for two hours at room temperature. After three washes with the Washing Buffer, the samples were mounted in ProLong Diamond Antifade Mountant (Invitrogen, P36970) with #1 coverglass (VWR, 48393-106).

Primary antibodies used were: rat anti-SOX2 (1:300, Invitrogen, 14-9811-82), rabbit anti-NKX2-1 (1:300, Abcam, 76013), rabbit anti-cleaved Caspase 3 (1:300, Cell Signaling Technology, 9661), rabbit anti-phospho-Histone H3 (1:300, Cell Signaling Technology, 3377), rabbit anti phospho-MLC2 (1:100, Cell Signaling Technology, 3674), rabbit anti-mCherry (1:500, Abcam, 167453, for labeling tdTomato), biotinylated Hyaluronic Acid Binding Protein (1:200, Sigma-Aldrich, 385911), mouse anti-GM130 (1:150, BD Biosciences, 610822), rabbit anti-phospho-FAK (1:200, Invitrogen, 700255), goat anti-NKX6-1 (10 µg/mL, R&D Systems, AF5857), rabbit anti-TBX1 (1:50, Invitrogen, 34-9800), mouse anti-ISL1 (1:30, DSHB, 40.2D6), mouse anti-ALDH1A2 (1:50, Santa Cruz Biotechnology, sc-393204), goat anti-SHH (1:100, R&D Systems, AF464). Dye-conjugated secondary antibodies were from Jackson ImmunoResearch (with Alexa Fluor 647, Cy3, or Alexa Fluor 488) and used at 1:300 dilution, except the streptavidin-Alexa Fluor 647 for HABP detection (1:300, Invitrogen, S32357).

Tissue sections were imaged with a Nikon Ti inverted microscope with a W1 spinning disk scanner (Yokogawa, CSU-W1) using a 20x or 40x objective (Nikon, MRH01401). Z-stacks with 0.3-0.9 µm step size were acquired and maximum intensity projection was performed with a custom Python program.

### Hybridization Chain Reaction (HCR)

Probes for HCR fluorescent in situ hybridization were designed by the insitu_probe_generator (https://github.com/rwnull/insitu_probe_generator) ^56^ using the coding sequences of chick *PTCH1*, mouse *Ptch1*, and mouse *Hhip* from the Ensembl Release 112. Generated probes were blasted against the respective species’ transcriptome to exclude potential off-target probes. The designed probes with HCR 3.0 barcode sequences (Molecular Instruments) were synthesized as 50 pmol oligo pools (oPools) by Integrated DNA Technologies and reconstituted in the TE buffer (Qiagen, 19086) to 1 µM.

Dissected foregut tissues were fixed in a 1.7 mL microcentrifuge tube with 4% PFA for one hour at room temperature, and washed twice with PBS (5 minutes each). The samples were then dehydrated and permeabilized with 70% ethanol (VWR, V1001) in PBS for one hour at room temperature. The samples were equilibrated with the probe wash buffer (Molecular Instruments) for 10 minutes at room temperature and then with the probe hybridization buffer (Molecular Instruments) for 30 minutes at 37°C (without rocking). Probes were diluted to 40 nM in 100 µL of hybridization buffer, incubating the samples overnight at 37°C without rocking. Labeled samples were washed twice with the wash buffer, then twice with 5X SSCT (5X Saline-Sodium Citrate buffer [Invitrogen, 15557044] and 0.1% Tween 20 [Sigma-Aldrich, P9416] diluted in PBS), and finally with the HCR amplification buffer (Molecular Instruments), 20 minutes each at room temperature. The HCR amplifiers with Alexa Fluor 647 or Alexa Fluor 546 (Molecular Instruments) were denatured at 95°C for 90 seconds and annealed at room temperature for 30 minutes. The amplifiers were diluted in the amplification buffer at 37 nM per strand, which was incubated with the sample overnight at room temperature without rocking. Two 5X SSCT washes were performed before embedding and sectioning in O.C.T. A 30-minute DAPI labeling in the Blocking Buffer was performed on the sections before mounting.

### Single-cell RNA sequencing (scRNA-seq)

#### Library preparation and sequencing

Control E3.5 (N = 6), E4.0 (N = 4), and cyclopamine-treated E4.0 (N = 9) foreguts were dissected and dissociated to single cells with TrypLE (Gibco, 12604013) at 37°C with intermittent trituration with a 25 Gauge needle. Dissociated cells were spun down at 800 g for 3 minutes at 4°C and washed twice with cold PBS. We performed MULTI-seq lipid barcode labeling on ice to multiplex the samples as described (Sigma-Aldrich, LMO001)^57^. After rinsing off the unlabeled barcode and anchors, the cell suspension was passed through a 40 µm Falcon cell strainer (VWR, 21008-949), spun at 800 g for 5 minutes at 4°C, and resuspended to 20 µL at a density of ∼500 cells/µL. All cells were used for Gel Bead-In Emulsions (GEM) with a Chromium Controller (10x Genomics) per manufacturer’s instructions. Library construction was performed with the Chromium Next GEM Single Cell 3’ Kit (v3.1) with dual index (10x Genomics). Quality control of the library was performed by the Biopolymers Facility at Harvard Medical School.

The 10-nM library was sequenced with NovaSeq 6000 platform at the Biopolymers Facility at Harvard Medical School, with the configuration of 28 bp for cell barcode 1 and UMI, 8 bp for i7 index, 10 bp for i5 index, and 90 bp for the transcript.

#### Demultiplexing by embryo and sample origin (MULTI-seq)

Samples of different groups were demultiplexed using the deMULTIplex package in RStudio (https://github.com/chris-mcginnis-ucsf/MULTI-seq) based on the counts of MULTI-seq barcodes. Leveraging the intrinsic genetic variation of chick embryos, individual cells were assigned to different embryos within each sample using the vireo package (https://github.com/single-cell-genetics/vireo)^58^. Doublets identified from MULTI-seq or vireo analysis were excluded from downstream analysis.

#### Quality control and clustering

Single-cell data was processed with Seurat V4 in RStudio (https://github.com/satijalab/seurat)^59^. A set of criteria were used to select high quality cells for downstream analysis: nCount_RNA >= 1000, nFeature_RNA >= 500, log10GenesPerUMI > 0.80, percent.mito < 0.18, percent.rbc < 0.20. After filtration, we ended up with 8,021 cells for control E3.5, 4,915 cells for control E4.0, and 4,829 cells for cyclopamine-treated E4.0. Differently treated samples were integrated using SCTransform of Seurat after regression of cell cycle and mitochondrial fractions. We used Uniform Manifold Approximation and Projection (UMAP) to perform dimension reduction of the integrated data. Louvain algorithm was used to cluster cells with 40 principal components based on the elbow plot, with a resolution of 0.6, resulting in 15 clusters. We annotated the clusters by their marker genes identified by FindMarkers, and verified their spatial distribution by HCR or immunostaining. For each cell cluster, we computed their abundance in total cells from the same embryo inferred by demultiplexing.

#### Differentially expressed gene (DEG) analysis

Given the demultiplexed biological replicates in each group, we performed pseudo-bulk DEG analysis using the glmGamPoi package in RStudio (https://github.com/const-ae/glmGamPoi)^60^. Cells of the dorsal sub-epithelial mesenchyme cluster were grouped into control (E3.5 and E4.0) and cyclopamine. Genes with adjusted *P* values < 0.001 were identified as DEGs. Volcano plot was generated by the EnhancedVolcano package (https://github.com/kevinblighe/EnhancedVolcano). Control group-enriched DEGs were imported into Shiny GO V0.77 (http://bioinformatics.sdstate.edu/go/)^61^ using the AmiGo 2 gene ontology database (https://amigo.geneontology.org/amigo) with a false discovery rate threshold of 0.1.

For ACTA2 expression analysis, as this gene is expressed at very low level at E4, prior to smooth muscle differentiation, we performed DEG analysis with FindMarkers in Seurat.

#### Cell-cell communication analysis

LIANA framework (https://github.com/saezlab/liana-py)^62,63^ was used to analyze potential ligand-receptor communication pathways between the dorsal epithelium and the dorsal mesenchymal cell types. CellPhoneDB^64^ with 100 permutations was used to identify ligand-receptor pairs, and those with *P* values < 0.05 were selected.

### Image analysis of immunofluorescence tissue sections

#### SOX2 gradient

SOX2 intensities were quantified in 12 equal, dorsal-ventrally spaced bins along the foregut epithelium in Fiji, and normalized to the most dorsal bin.

#### pMLC apical-basal ratio

Line profiles normal to the epithelial surface were drawn in Fiji, and the local peak intensity values at the apical and the basal side of the epithelium were taken. The ratio between the apical and basal intensity of the same tissue was calculated, which reflects the extent of apical constriction.

#### Cell roundness

MLC2-tdTomato-electroporated cells were manually segmented and cell roundness was calculated in Fiji (https://imagej.net/ij/docs/menus/analyze.html) by:

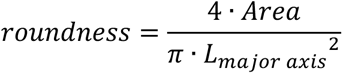

#### Cell orientation analysis

Relative positions of the Golgi and the nucleus were used to define cell orientation. Nuclei were segmented with StarDist in TrackMate. The Golgi apparatus was segmented with the LoG detector in TrackMate. The coordinates of the segmented structures were exported to MATLAB, and we wrote a custom code to pair the Golgi and the nucleus with the costMatrix function. Due to the thinness of sections, we often had extra, unpaired nuclei. We thus defined the nucleus-Golgi vectors and computed the angular histograms of their directions. For the epithelium-mesenchyme interface, 0 degree was defined as the vector pointing normal to the epithelium. For bead implantation, 0 degree was defined as the vector pointing towards the center of the bead.

#### pHH3 and CC3 analysis

As pHH3 and CC3 labels are sparse within the tissue, we define their frequencies by the number of positive cells per unit area of the mesenchyme. Positive cells were counted manually and the area of mesenchyme was measured in Fiji.

#### Epithelium right-left distance ratio with one-sided electroporation

We defined the midline of the epithelium by connecting the dorsal and ventral points with the highest curvature. The one-sided distance was measured as the perpendicular line segment from the center of the electroporated cell patch’s basal surface to the midline. The ratio of right (electroporated) to left (non-electroporated) distances was then calculated.

### Morphometric analysis of the foregut epithelium

The time-lapse videos of TES in slice culture were first processed in Fiji, and the Segment Anything Model 2 (SAM 2, https://github.com/facebookresearch/sam2)^65^ was applied to segment the epithelium as a mask. The mask was then processed with the Open Source Computer Vision Library (OpenCV, https://github.com/opencv/opencv) to extract the contour and conduct downstream morphometric analyses.

#### Video pre-processing

The Bleach Correction function in Fiji^66^ was used to correct for photobleaching (Supplementary Fig. 3g), and StackReg to correct translational and rotational drifts in the rigid body mode, given that the morphological change of the slice was small between two consecutive frames. The corrected video was then cropped to center the epithelium and Gaussian blurred to reduce cellular granularity for higher segmentation accuracy.

#### Automated segmentation by SAM 2

The pre-trained sam2.1_hiera_large model of SAM 2 was used to segment the foregut epithelium in the time-lapse video. At least one point on the epithelial region was selected as guide. However, when a single point selection led to misidentification, additional points were used to refine the segmentation. In some cases, bounding boxes were applied in conjunction with points to further constrain the region and isolate the epithelium accurately. This combination of point selection and bounding boxes ensured precise segmentation yielding a > 95% success rate for all chick videos. For the remaining frames, we manually segmented the epithelium in Fiji and replaced the wrong segmentations. The segmentation results were saved as binary masks.

However, segmentation of mouse videos remained challenging because the nTnG mouse slices had nuclear labeling, which made the epithelium more granular than the GFP chick slices. Therefore, in this work we only quantify the morphological evolution of chick TES.

#### Morphometric analysis

From the binary segmentation, we found the outline of the shape using the OpenCV wrapper in Python. For each frame of a given sample, we fitted an ellipse using the fitEllipse function in OpenCV. The height and width of the rectangle in which the ellipse is inscribed was used as measures of the shape’s dimensions. We chose to fit an ellipse because we found it yields a more representative measure of the shape than, for example, a rectangle enclosing the 2D point set. Additionally, we computed the arc length and area of the shape using OpenCV.

To calculate the curvature of an outline, we first fitted a B-spline to the 2D point set of each frame in a given video. This was done using the function BSplineFunction in Mathematica. The degree of the B-spline was chosen to be one-quarter of the total number of points in the outline for a given frame. We found this value to be a good compromise between smoothing the noisy data sufficiently for the following operations (namely, computing the curvature) and preserving the high-curvature areas in the neck region of interest. For example, the area and arc length of the fitted curve agree well with the values obtained from the discrete points using OpenCV described above. This thus resulted in a smooth curve *γ*(*s*), with arc-length parametrization *s*. From this, the curvature was computed using the equation:

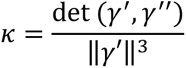

To define the neck region, we proceeded as follows: For each frame, we first selected the point *s*=*s_0_* at which the curvature reaches its global minimum. We then selected a second point *s_1_* from the region *s*=*s_0_*+0.5±0.2 at which the curvature takes its local minimum in this region. We followed this procedure instead of simply picking the two smallest-curvature points to reduce errors due to noise and segmentation inaccuracies. We found that this method is robust, and the points identified in this way correspond well with those identified manually. Overall, across all videos, the procedure only failed in a few isolated frames when segmentation issues occur. The two identified points then define the neck. We repeated this procedure for all frames of a given video, resulting in a time evolution of the neck curvatures. To mitigate the impact of segmentation errors and noise, we performed a moving median over three frames for the two neck curvatures. The curvature was normalized by multiplication with the initial neck width of the epithelium to account for size variation in different samples.

To average the quantities from several videos of a specific developmental stage, we first interpolated the data obtained from each video. This is necessary since the videos were recorded at different temporal resolutions. Before interpolating, we removed outliers in the data, defined as points that deviate more than 2.5 standard deviations from the average of a given video. This helps reduce the effect of segmentation errors. We then aligned the videos in time using the manually identified frame at which the transition occurs. We computed the median and median deviation (MAD) across all videos for a given point in time and repeated this for all time points to minimize the effect of outliers due to inaccurate segmentation or contour extraction. For experimental groups with N < 6, individual curves were plotted instead of the population average.

#### Image central moments

We chose to use the central moments for extracting morphological information from the binary epithelial masks^67^. Central moments were calculated as:

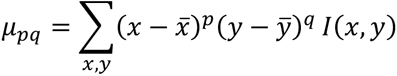

Here 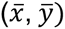 is the mask centroid, *I*(*x, y*) represents the pixel intensity (1 or 0) at (*x, y*). The second-order central moments were normalized as:

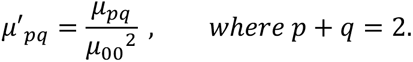

### PIV analysis

Drift-corrected, contrast-enhanced time-lapse videos were imported to PIVlab V2.62 in MATLAB^68^. The image frames were first pre-processed to highlight the individual cells, with CLAHE with a window size of 25 pixels, highpass filtering with a kernel size of 35 pixels, and Wiener2 denoising with a window size of 5 pixels. The PIV settings used were: FFT window deformation algorithm, Pass 1 with 128-pixel integration area and 64-pixel step size, Pass 2 with 64-pixel integration area and 32-pixel step size, and high correlation robustness. The calculated velocity field was filtered by X and Y velocities and a correlation coefficient filter of 0.4 to exclude outliers. The velocity fields were then temporally averaged with a window of one hour. Velocity vectors were recolored by their directions using Cyclic Color Map in MATLAB (by Chad Greene, https://www.mathworks.com/matlabcentral/fileexchange/57020-cyclic-color-map).

The strain rate and the divergence were computed based on the gradients of the averaged velocity fields with an additional 2X2 spatial binning using a custom Python code. We also calculated the eigenvectors of the two-dimensional strain rate tensor to indicate the principal direction of strain.

### Single-cell tracking and trajectory analysis

Cells were tracked in Fiji TrackMate using the StarDist detector without frame gaps^69^. The trajectories and average velocity vectors were plotted with a custom MATLAB code. The angular histograms were computed with the velocity vectors in MATLAB. Mean squared displacements (MSD) were plotted for each trajectory (starting time defined as 0) with a population average and SD. The average *MSD-t* curve was fitted by a power function describing anomalous diffusion^70^:

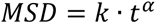

Here, *α* = 1 indicates normal diffusion, whereas *α* = 2 indicates ballistic motion. 1 < *α* < 2 suggests super-diffusion which can be due to active cell migration in a crowded environment.

We also quantified the cell shape in GFP chick samples with good image contrasts. Similarly, cells were tracked with StarDist in TrackMate, and the aspect ratio (defined as the inverse of cell roundness) was calculated and plotted for individual cells.

For tracking cells in 3D, we took live imaging stacks with 2-µm z step size (8-9 z planes in total). The image stack was corrected for translational drift using Correct 3D drift^71^ (https://imagej.net/plugins/correct-3d-drift). Cells were segmented in 3D using a custom Arivis pipeline including background subtraction, denoising, and the Blob Finder segmenter. The segmented cells were then tracked in Arivis using the nearest neighbor algorithm with a maximal inter-frame displacement of 5 µm. The cell tracks were then exported to MATLAB for plotting and angle calculations.

### Statistics and reproducibility

The number of samples used for each experiment and statistical tests were indicated in the figure legends. The sample sizes were not pre-determined. GraphPad Prism software was used to plot the data.

## Data availability

The chick single-cell RNA-sequencing data will be deposited to the Gene Expression Omnibus of NCBI. For inquiries about other data or materials used in this study, please reach out to the corresponding authors.

## Code availability

The custom codes used for image processing and data analysis will be uploaded to GitHub prior to the publication date.

---

**Supplementary Figure 1.**
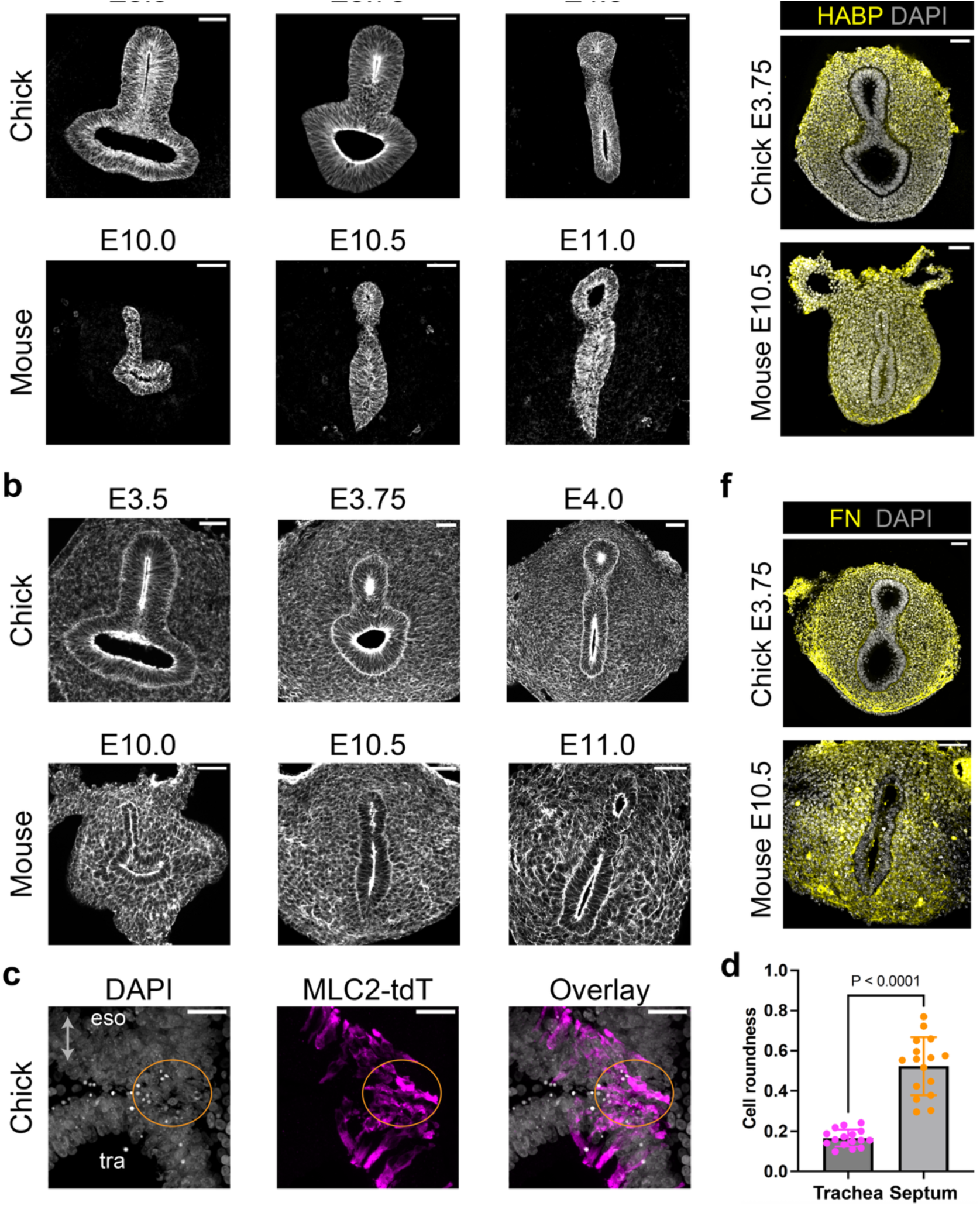
Morphological comparison of chick and mouse foregut cells during TES. **(a)** Immunofluorescence of E-cadherin in transverse sections of chick and mouse foreguts. **(b)** Phalloidin staining of F-actin in transverse sections of chick and mouse foreguts. **(c)** High-resolution immunofluorescence image of electroporated MLC2-tdTomato in the foregut epithelium, showing that the septal cells (orange ellipse) are less elongated than the cells in the esophagus (eso) or the trachea (tra). **(d)** Quantification of cell roundness in electroporated septal and tracheal cells in (c) (N = 16 cells from 4 biological replicates). **(e)** Hyaluronic acid binding protein (HABP) staining of chick and mouse TES. **(f)** Immunofluorescence of fibronectin (FN) in chick and mouse TES. Scale bars: 50 µm (a,b,e,f), 20 µm (c). Error bars: SD.

**Supplementary Figure 2.**
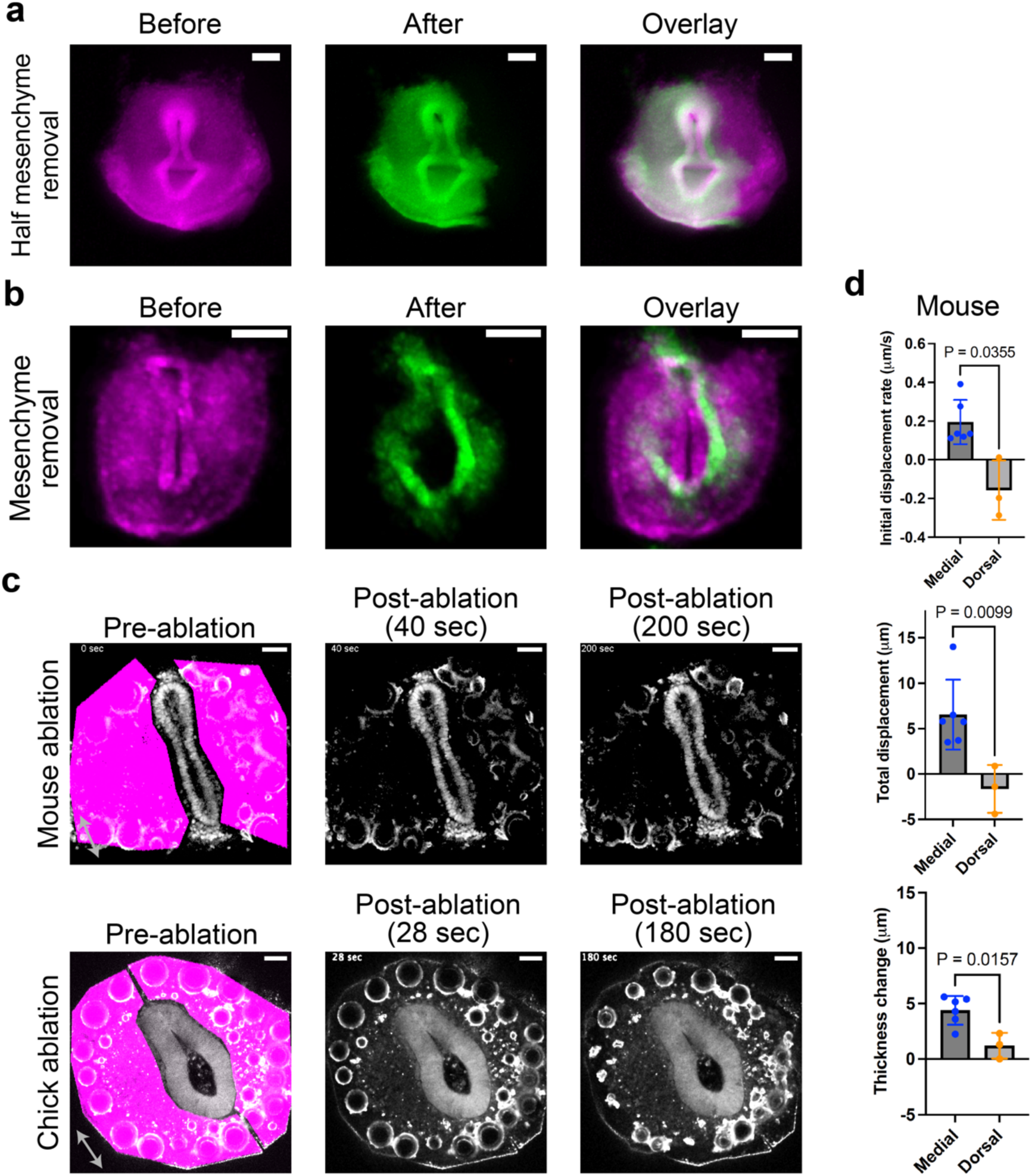
Constrictive mesenchymal force deforms the epithelium during TES. **(a)** Stereoscope imaging of an E3.75 chick foregut slice before (magenta) and after (green) one-sided mesenchyme removal. **(b)** Stereoscope imaging of the transverse slice of an E10.5 nTnG mouse foregut before (magenta) and after (green) surgical removal of the mesenchyme. **(c)** Live imaging of an E10.5 nTnG mouse slice (top) and an E3.75 GFP chick slice (bottom) after two-photon laser ablation of all surrounding mesenchyme (magenta, ∼10-µm ablation depth in z axis). **(d)** Quantifications of mouse dorsal and medial epithelial dynamics after ablation as specified in Fig. 2c. N = 3 or 6 samples. Scale bars: 100 µm (a,b), 50 µm (c). Error bars: SD. P values: Two-tailed Welch’s t test.

**Supplementary Figure 3.**
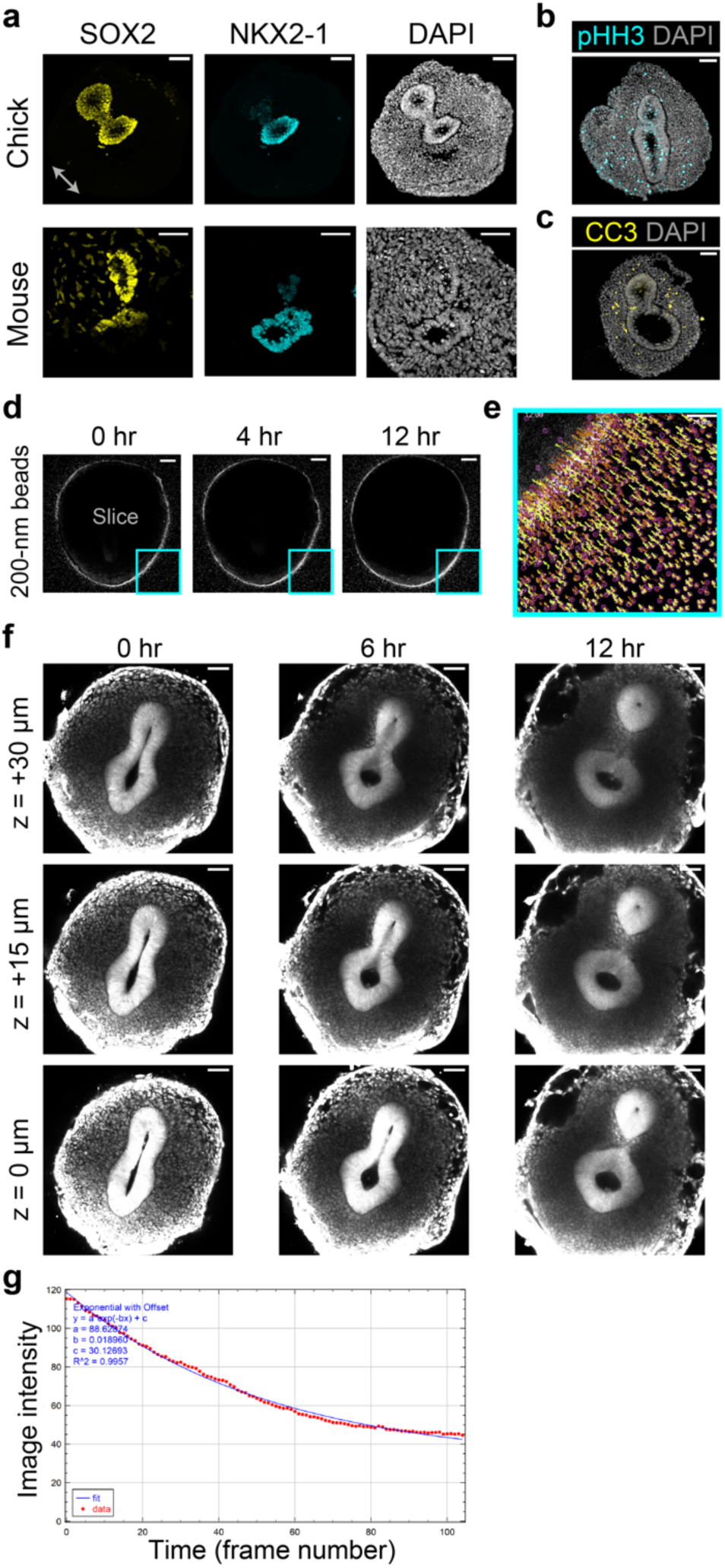
Ex vivo slice culture preserves tissue patterning and cell viability, recapitulating TES in the chick and the mouse. **(a)** Immunofluorescence of SOX2 and NKX2-1 in transverse sections of chick and mouse slices cultured for 24 hours. **(b,c)** Immunofluorescence of phospho-Histone H3 (b) and cleaved Caspase 3 (c) in transverse sections of chick slices cultured for 24 hours. **(d)** Live imaging of 2-nm red fluorescent beads in collagen gel with a slice sample in the middle. **(e)** Single-particle trajectories of the beads in the cyan box in (d) for over 12 hours indicate the stability of x, y, and z positions of the setup during live imaging. **(f)** Live imaging of an E4.0 GFP chick slice at different z depths. **(g)** Live imaging intensity decay over time (blue) overlaid by an exponential decay model (red). Scale bars: 50 µm.

**Supplementary Figure 4.**
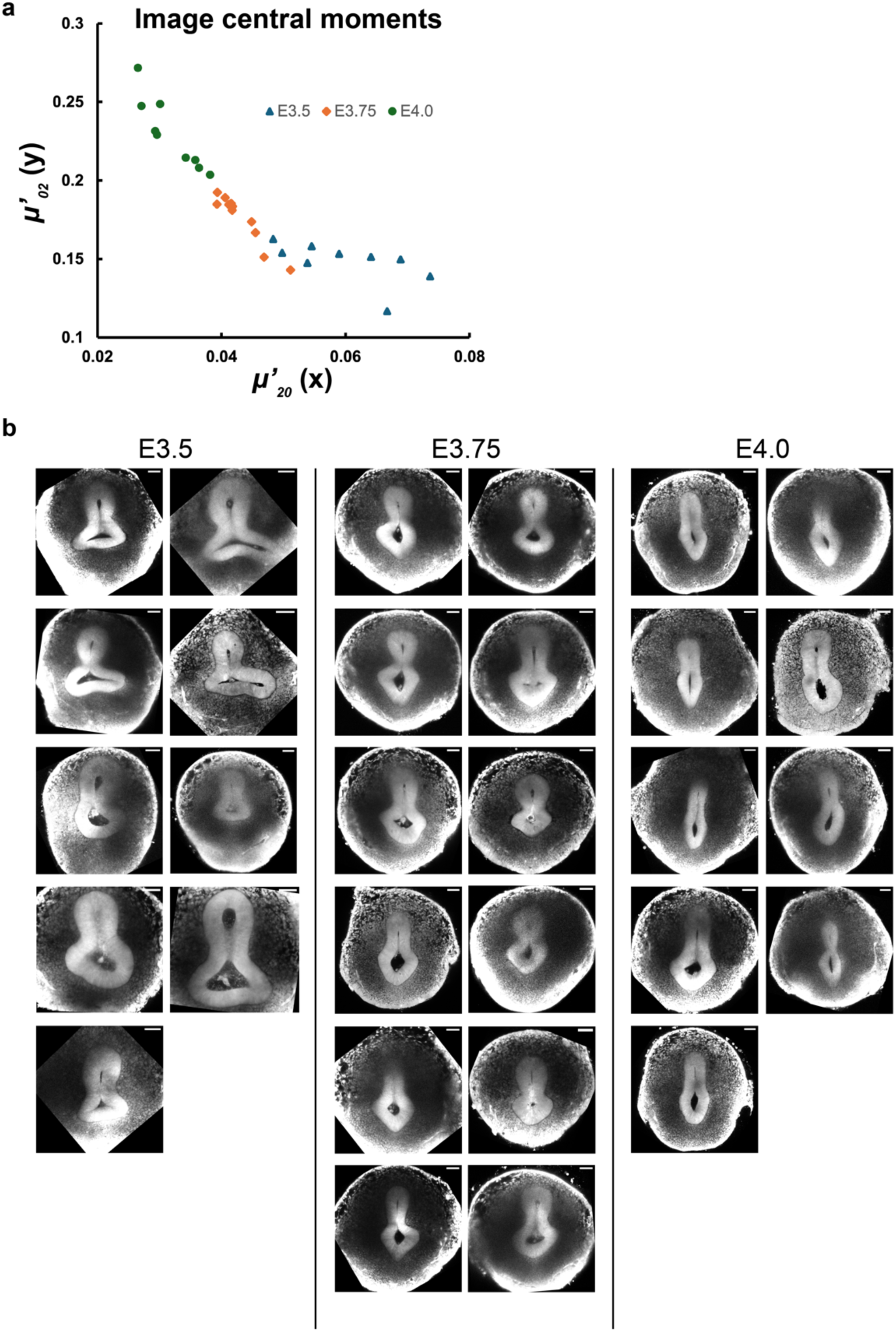
Intragroup and intergroup variations of the slice culture. **(a)** Normalized second-order central moments of the segmented chick E3.5, E3.75, and E4.0 epithelial images indicate effective separation between groups (see Methods). **(b)** Raw images at time 0 (septum formation) used for segmentation and morphological quantifications in Fig. 3. Scale bars: 50 µm.

**Supplementary Figure 5.**
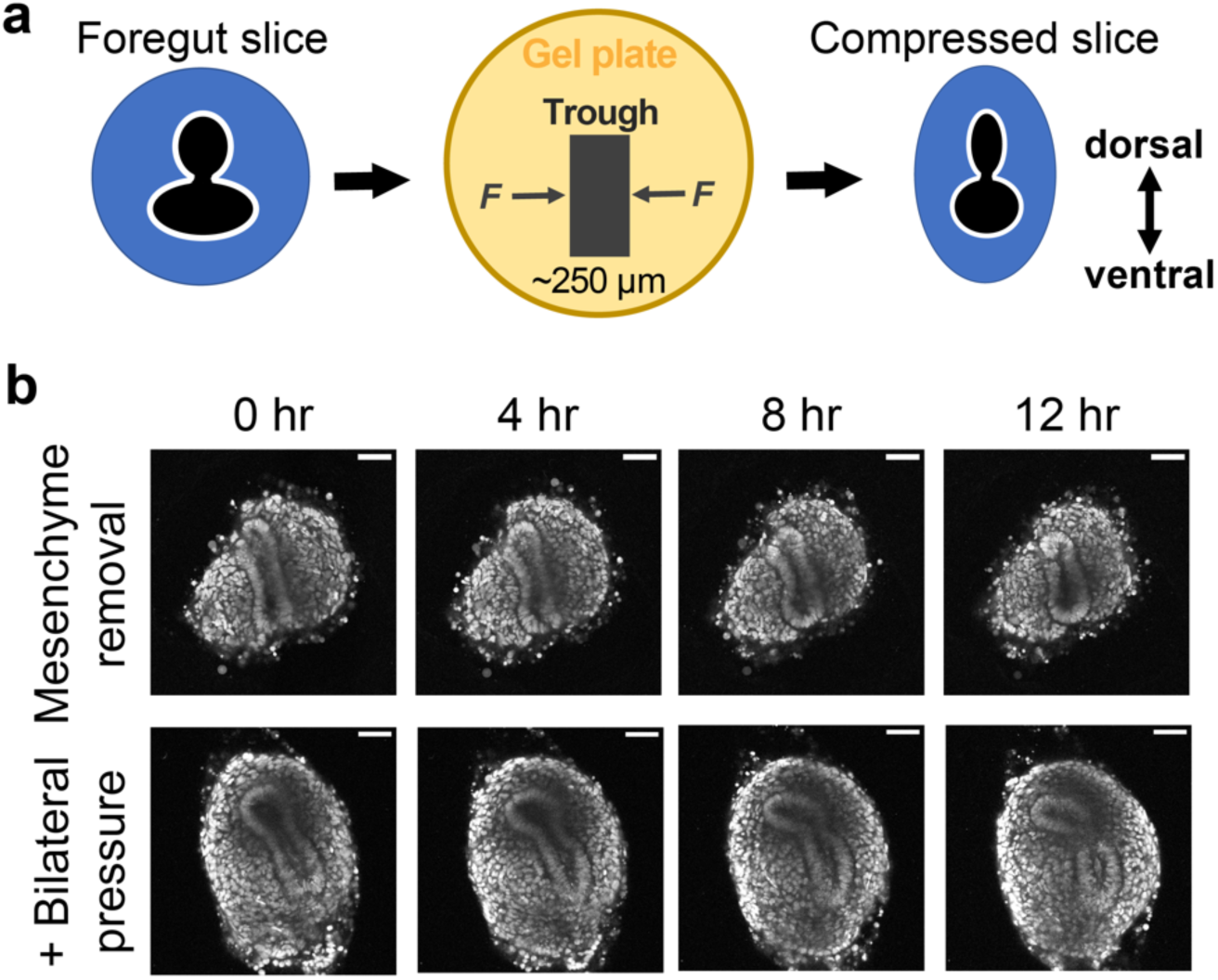
Bilateral compression of the slice culture. **(a)** Schematics of applying bilateral compressive pressure to the foregut slice culture by fitting the slice into a trough in collagen gel. Continued mesenchymal cell proliferation in confinement generates bilateral pressure on the epithelium. **(b)** Live imaging of E10.5 nTnG mouse slices with surgical removal of the mesenchyme, in the absence (top) or the presence (bottom) of bilateral pressure by a collagen gel trough. Scale bars: 50 µm.

**Supplementary Figure 6.**
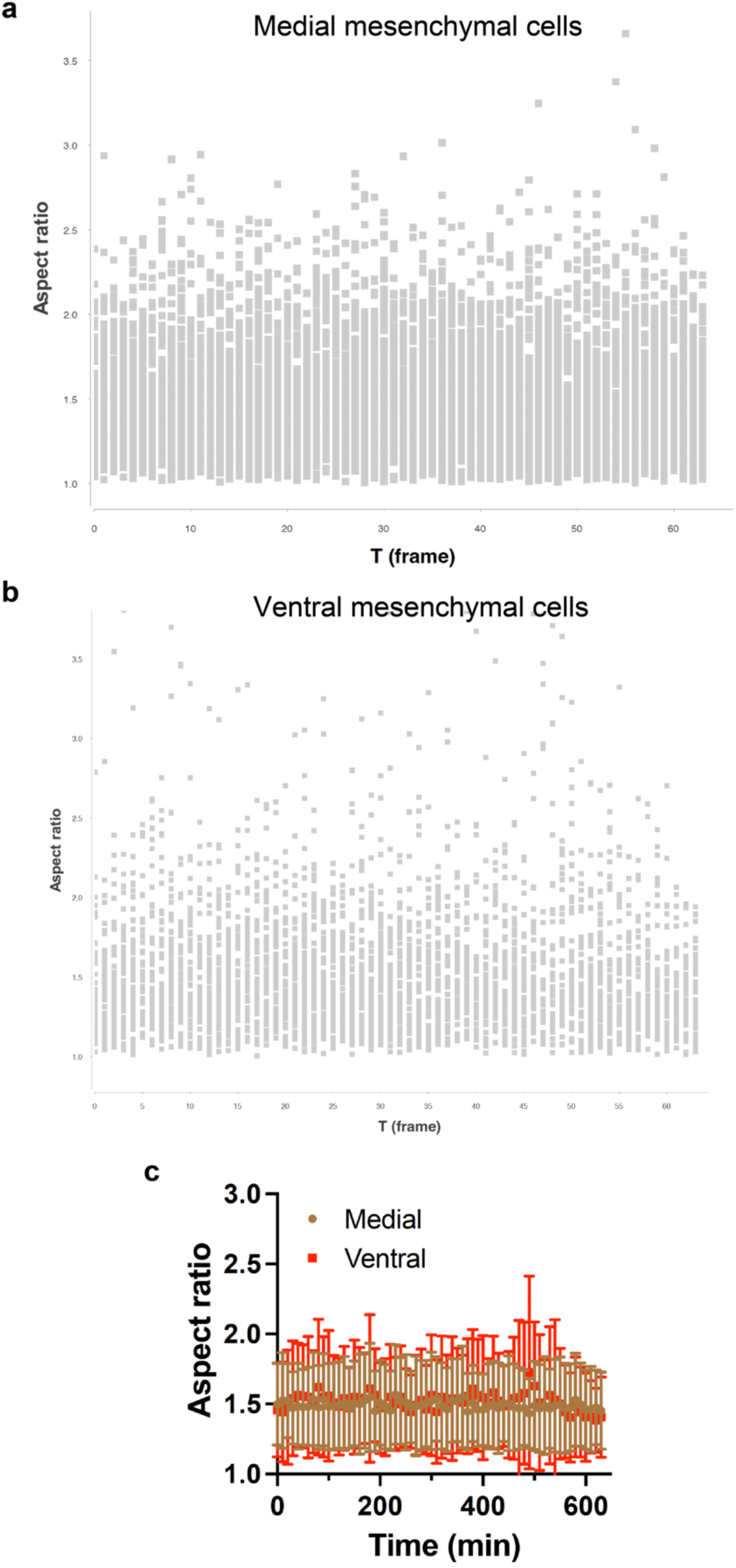
Mesenchymal cell shape evolution during TES. (a,b) Quantifications of medial (a) and ventral (b) mesenchymal cell aspect ratios from the chick E3.75 sample in Fig. 3b. **(c)** Populational averages of (a,b). Scale bars: 50 µm. Error bars: SD.

**Supplementary Figure 7.**
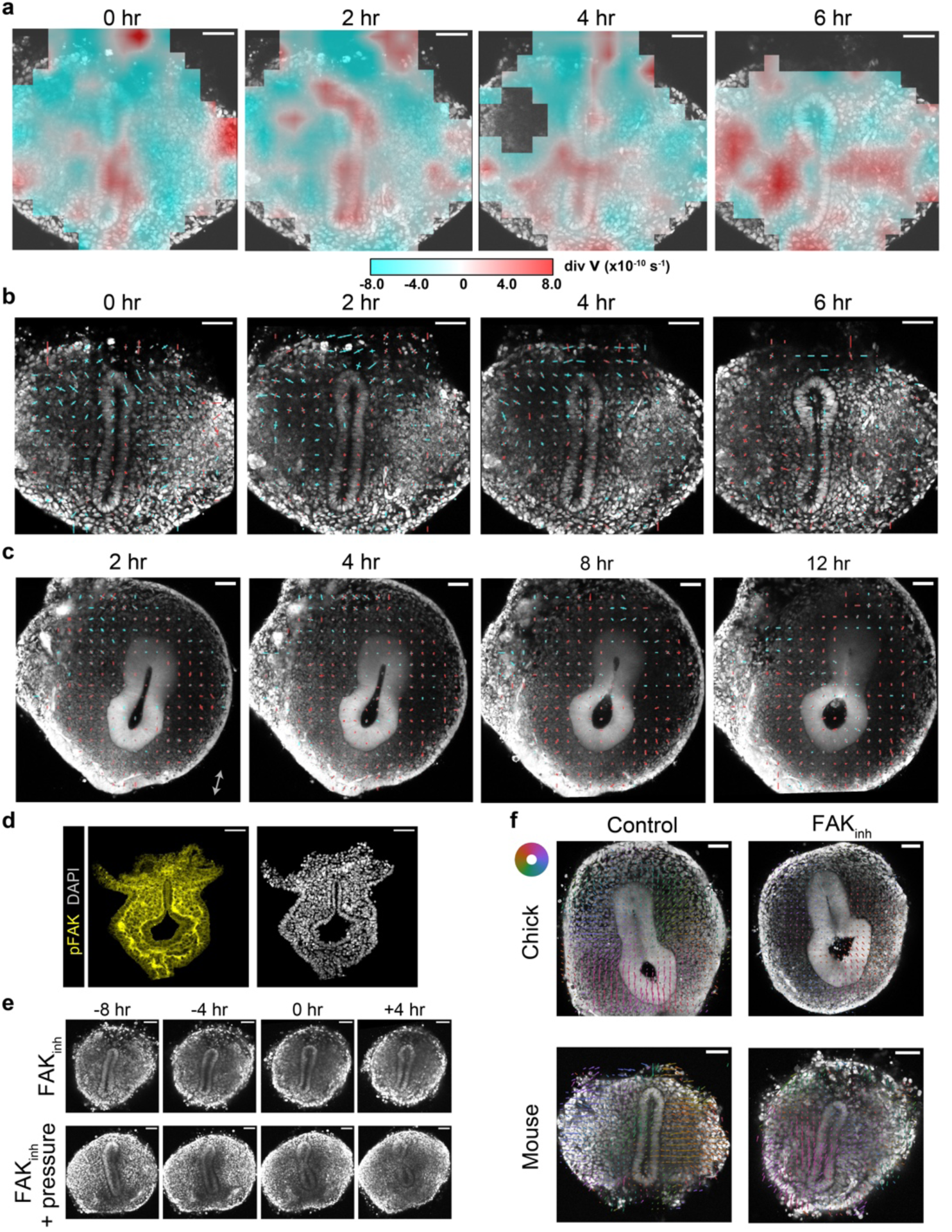
Directional mesenchymal flow contributes to the mesenchymal force and is essential for TES. **(a)** Mapping of the divergence of the velocity field from Fig. 5a. Cyan indicates negative divergence and red indicates positive divergence. **(b)** Principal strain rates calculated from the velocity field in Fig. 5a. Line segments are plotted along the direction of the eigenvectors of the strain rate tensor, and their lengths are proportional to the corresponding eigenvalues. Cyan indicates compressive strain and red indicates expansive strain. **(c)** Principal strain rates of an E3.75 GFP chick foregut slice culture. **(d)** Immunofluorescence of phospho-FAK in an E10.0 mouse section. **(e)** Live imaging of E10.0 nTnG mouse slices with 2.5 µM FAK inhibitor, in the absence (top) or the presence (bottom) of bilateral pressure by a collagen gel trough. **(f)** PIV analysis of control and FAK inhibitor-treated chick and mouse slices. The color wheel indicates the directions of PIV vectors. Scale bars: 50 µm.

**Supplementary Figure 8.**
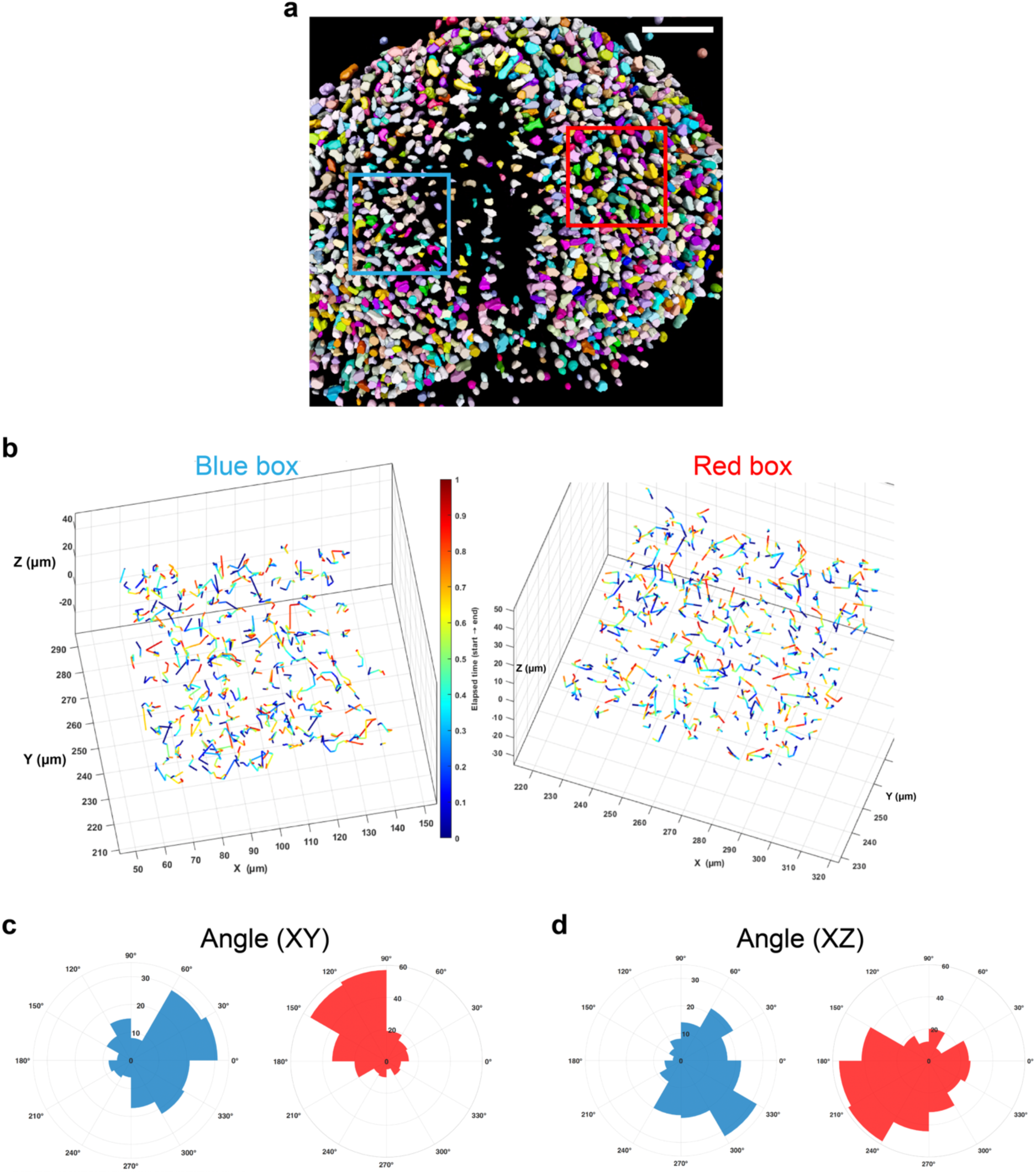
3D tracking of cells confirms the convergent mesenchymal movements. **(a)** 3D segmentation of cells from the data in Fig. 5b (see Methods). Cell colors are randomly assigned. **(b)** Individual cell trajectories in the blue and red boxes in (a). Trajectories are color-coded by time. **(c,d)** Angular histograms of cell displacement vectors from (b) in xy (c) and xz (d) planes. Scale bar: 50 µm.

**Supplementary Figure 9.**
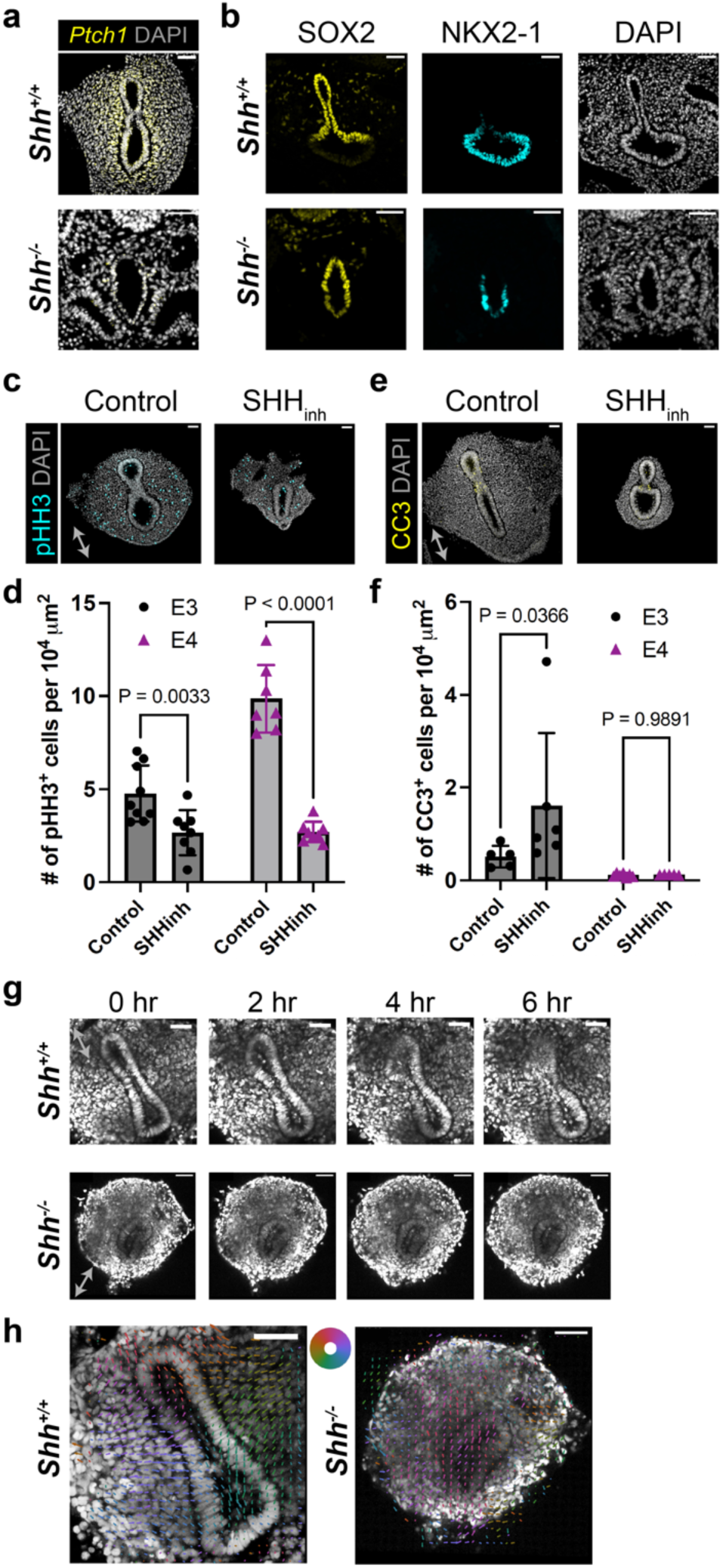
SHH signaling is essential for generating the convergent mesenchymal force. **(a)** HCR-FISH of *Ptch1* in transverse sections of E10.5 *Shh^+/+^* or *Shh^-/-^*mouse foreguts. **(b)** Immunofluorescence of SOX2 and NKX2-1 in transverse sections of E10.5 *Shh^+/+^* or *Shh^-/-^* mouse foreguts. **(c-f)** Immunofluorescence of phospho-Histone H3 (c) and cleaved Caspase 3 (e) in transverse sections of E3.75 chick slices with or without *in ovo* SHH inhibitor treatment. Quantification of pHH3 (d) and CC3 (f) frequencies reveals a significant impact on cell proliferation by SHH inhibition. **(g,h)** Live imaging and PIV analysis of E10.5 *Shh^+/+^;nTnG/nTnG* or *Shh^-/-^;nTnG/nTnG* mouse slices. Scale bars: 50 µm. Error bars: SD. P values: Two-tailed t test.

**Supplementary Figure 10.**
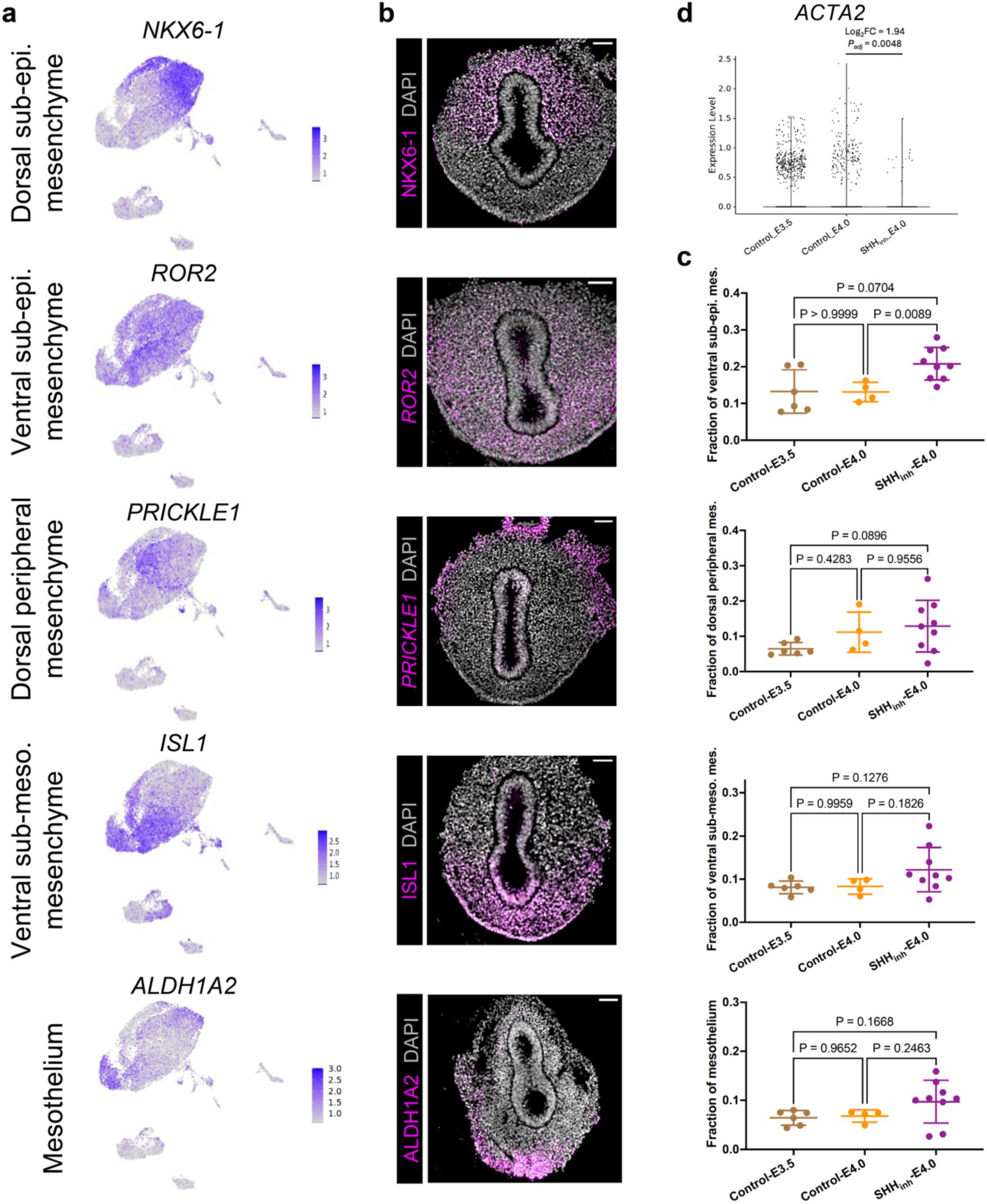
Single-cell RNA sequencing analysis of mesenchymal subtypes. **(a)** Seurat FeaturePlots of mesenchymal subtype marker genes. **(b)** Immunofluorescence (NKX6-1, ISL1, ALDH1A2) or HCR-FISH (*ROR2*, *PRICKLE1*) of mesenchymal subtype marker genes in E3.75 chick foregut sections. **(c)** Fractions of the mesenchymal cell subtypes in individual embryos recovered from scRNA-seq data. Each data point represents one embryo. P values are calculated by one-way ANOVA with Dunnett’s test. **(d)** Expression levels of ACTA2 (Smooth Muscle Actin). Adjusted P value is calculated in Seurat with Bonferroni correction. Scale bars: 50 µm.

**Supplementary Figure 11.**
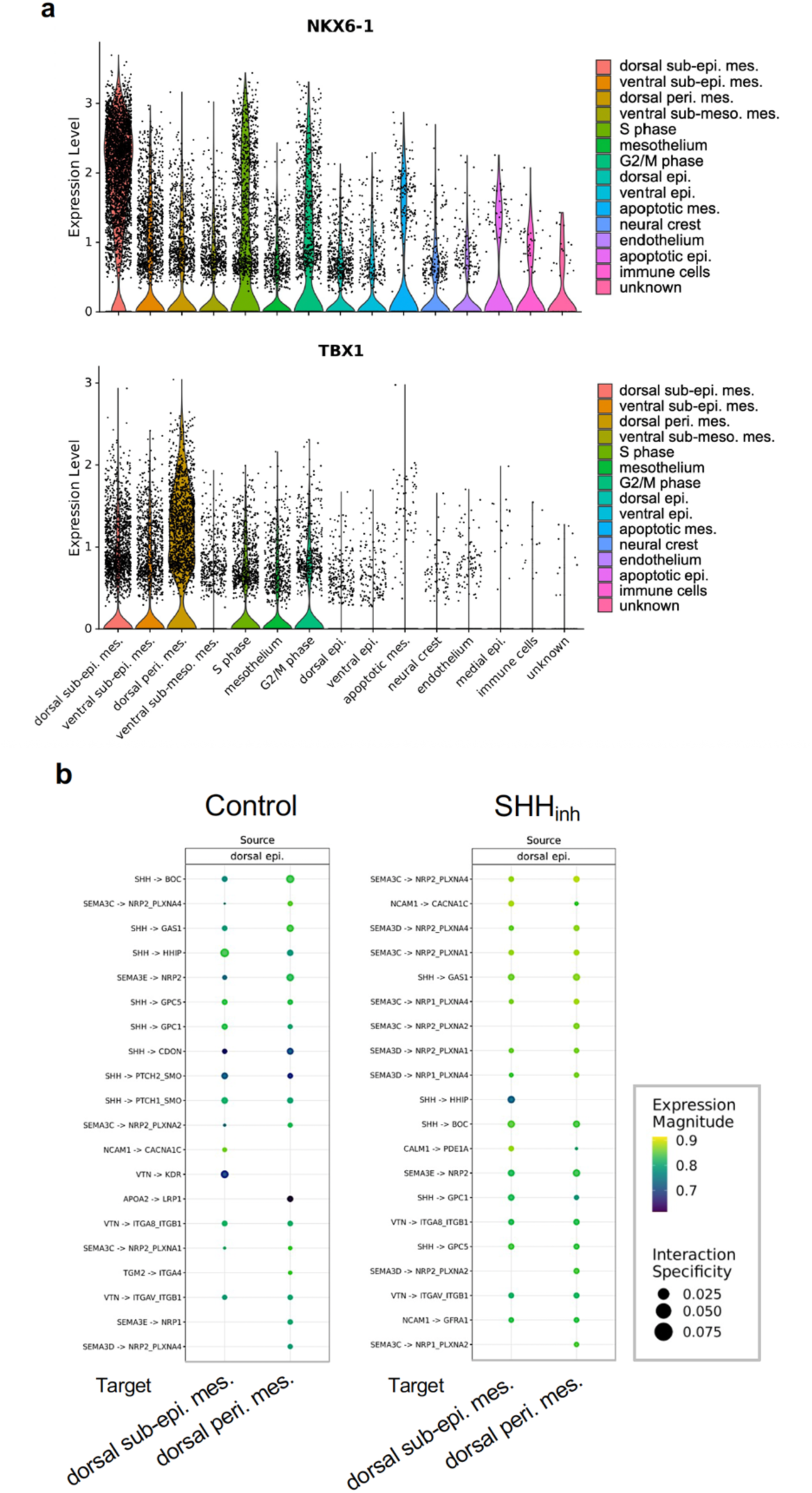
Single-cell RNA sequencing analysis of dorsal mesenchymal cell types. **(a)** Violin plots of NKX6-1 and TBX1 expression across different cell types. **(b)** LIANA cell-cell interaction analysis of possible communication pathways from the dorsal epithelium (source) to the dorsal sub-epithelial mesenchyme or the dorsal peripheral mesenchyme (target), in control (WT) and cyclopamine-treated (SHH_inh_) samples.

**Supplementary Figure 12.**
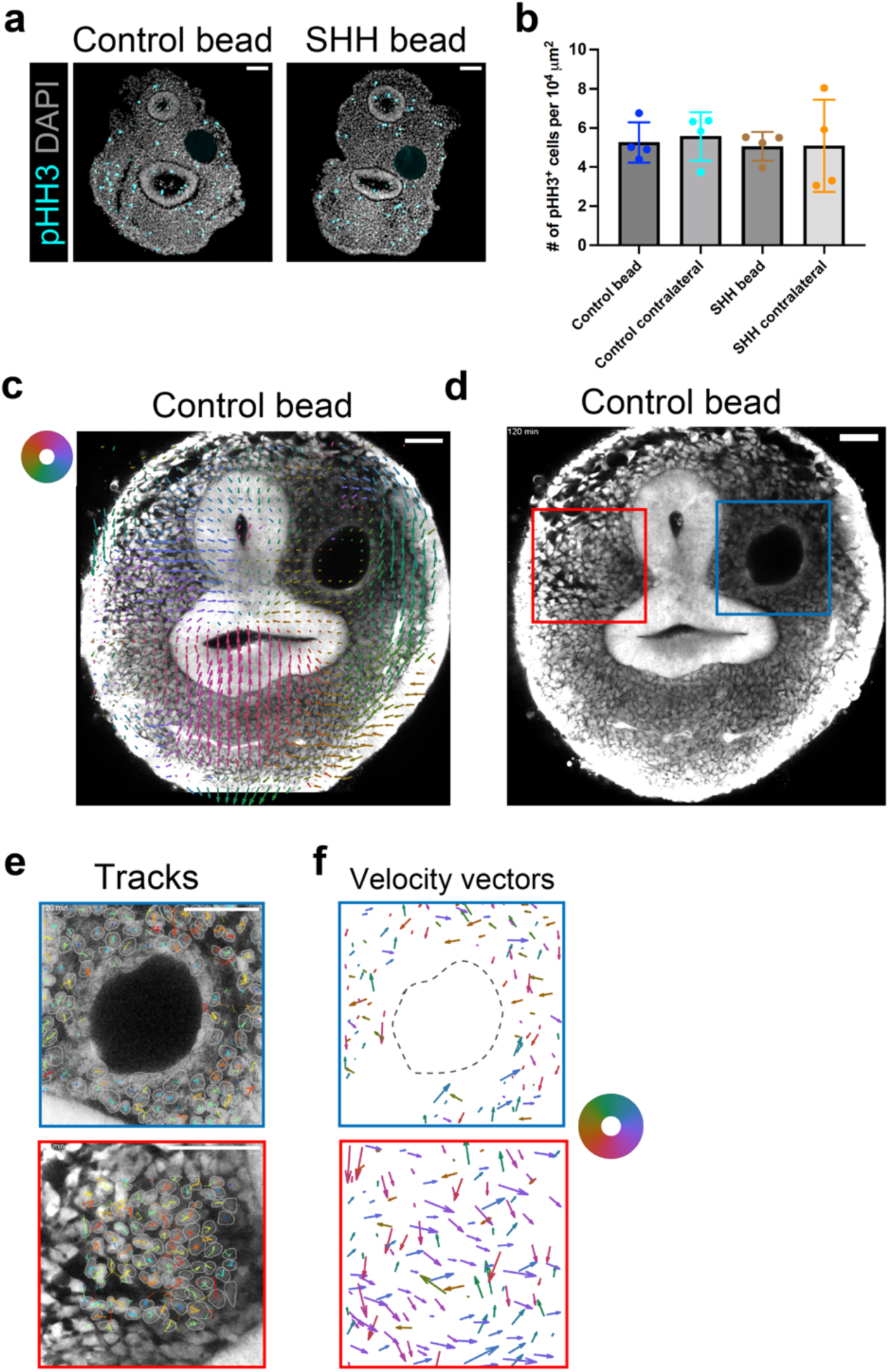
SHH bead does not induce mesenchymal over-proliferation. **(a)** Immunofluorescence of phospho-Histone H3 in E3.75 chick foregut slices 24 hours after implantation of a control or SHH-loaded bead. **(b)** Quantification of pHH3 frequencies near the implanted beads (N = 4 replicates). **(c)** PIV analysis of an E3.5 GFP chick slice with an implanted control bead. The color wheel indicates the directions of PIV vectors. **(d)** Live imaging of a SHH-bead-implanted GFP chick slice used for single-cell tracking. **(e,f)** Cell tracks (e) and average velocity vectors (f) of individual cell trajectories in the blue and red boxes of (d). Vectors are scaled with the speed and color-coded by their directions according to the color wheel. The dashed circle marks the bead. Scale bars: 50 µm. Error bars: SD.

## Description of supplementary movies

**Supplementary Movie 1.** 3D rendering of an E4 chick foregut immunostained by an E-cadherin antibody, related to Fig. 1a-c. Grid size: 200 µm.

**Supplementary Movie 2.** 3D rendering of an E10.5 mouse foregut immunostained by an E-cadherin antibody, related to Fig. 1a-c. Grid size: 200 µm.

**Supplementary Movie 3.** Tissue dynamics after laser ablation of the medial (left) or dorsal (right) sub-epithelial mesenchyme in an E3.75 GFP chick foregut slice, related to Fig. 2b-d. Scale bar: 50 µm.

**Supplementary Movie 4.** Tissue dynamics after laser ablation of all mesenchyme in an E10.5 nTnG mouse foregut slice, related to Supplementary Fig. 2c,d. Scale bar: 50 µm.

**Supplementary Movie 5.** Tissue dynamics after laser ablation of all mesenchyme in an E3.75 GFP chick foregut slice, related to Supplementary Fig. 2c,d. Scale bar: 50 µm.

**Supplementary Movie 6.** Slice culture of an E4.0 GFP chick foregut (left) and automatedly segmented epithelial contour (right), related to Fig. 3a-f. Orange line shows the segmented contour of the epithelium, and the red line is the spline fit of the contour for curvature calculation. Two local minima of curvature are marked by blue circles, and the connecting line segment indicates the neck width. Time is shown as HH:MM. Scale bar: 50 µm.

**Supplementary Movie 7.** Slice culture of an E10.5 nTnG mouse foregut, related to Fig. 3b. Time is shown as HH:MM. Scale bar: 50 µm.

**Supplementary Movie 8.** Slice culture with tracking of 200-nm red fluorescent beads, related to Supplementary Fig. 3d,e. Time is shown as HH:MM. Scale bar: 20 µm.

**Supplementary Movie 9.** Slice culture of an E3.75 GFP chick foregut after partial removal of the mesenchyme, related to Fig. 3g-i. Time is shown as HH:MM. Scale bar: 50 µm.

**Supplementary Movie 10.** Slice culture of an E3.75 GFP chick foregut after near-complete removal of the mesenchyme, related to Fig. 3g-i. Time is shown as HH:MM. Scale bar: 50 µm.

**Supplementary Movie 11.** Slice culture of an E3.75 GFP chick foregut after removal of the mesenchyme with bilateral compressive pressure, related to Fig. 3g-i. Time is shown as HH:MM. Scale bar: 50 µm.

**Supplementary Movie 12.** Slice culture of an E3.75 GFP chick foregut with 3 µM aphidicolin, related to Fig. 4d-f. Time is shown as HH:MM. Scale bar: 50 µm.

**Supplementary Movie 13.** Slice culture of an E10.5 nTnG mouse foregut used for PIV analysis, related to Fig. 5a. Time is shown as HH:MM. Scale bar: 50 µm.

**Supplementary Movie 14.** Slice culture of an E10.5 nTnG mouse foregut with 2D-tracked cell trajectories colored by average speeds, related to Fig. 5b-f. Time is shown as HH:MM. Scale bar: 50 µm.

**Supplementary Movie 15.** The same data as in Supplementary Movie 14 with randomly colored 3D cell segmentation (trajectories not shown for clarity), related to Supplementary Fig. 8.

**Supplementary Movie 16.** Slice culture of an E3.75 GFP chick foregut with 2.5 µM PF-573228, related to Fig. 5g-i. Time is shown as HH:MM. Scale bar: 50 µm.

**Supplementary Movie 17.** Slice culture of an E3.75 GFP chick foregut with 2.5 µM PF-573228 and bilateral compressive pressure, related to Fig. 5g-i. Time is shown as HH:MM. Scale bar: 50 µm.

**Supplementary Movie 18.** Slice culture of an E3.75 GFP chick foregut treated by cyclopamine

*in ovo*, related to Fig. 6g-i. Time is shown as HH:MM. Scale bar: 50 µm.

**Supplementary Movie 19.** Slice culture of an E3.75 GFP chick foregut treated by cyclopamine *in ovo*, with bilateral compressive pressure, related to Fig. 6g-i. Time is shown as HH:MM. Scale bar: 50 µm.

**Supplementary Movie 20.** Slice culture of an E3.75 GFP chick foregut after removal of the dorsal mesenchyme, related to Fig. 7h-j. Time is shown as HH:MM. Scale bar: 50 µm.

**Supplementary Movie 21.** Slice culture of an E3.75 GFP chick foregut after removal of the dorsal mesenchyme, with bilateral compressive pressure, related to Fig. 7h-j. Time is shown as HH:MM. Scale bar: 50 µm.

**Supplementary Movie 22.** Slice culture of an E3.75 GFP chick foregut implanted with a SHH-loaded bead, related to Fig. 8c-e. Time is shown as HH:MM. Scale bar: 50 µm.

**Supplementary Movie 23.** Slice culture of an E3.5 GFP chick foregut implanted with a SHH-loaded bead, related to Supplementary Fig. 12c-f. Scale bar: 50 µm.

## Notes

### Competing Interest Statement

The authors have declared no competing interest.

### Summary of Updates

We added analysis of single-cell trajectories in 2D and 3D, cell polarity analysis by the nucleus-Golgi vector, and control experiments of system stability and variations.

